# Sensing of dietary amino acids and regulation of calcium dynamics in adipose tissues through Adipokinetic hormone in *Drosophila*

**DOI:** 10.1101/2024.03.04.583442

**Authors:** Muhammad Ahmad, Shang Wu, Xuan Guo, Norbert Perrimon, Li He

**Author notes:** These authors contributed equally to this work (Muhammad Ahmad, Shang Wu).

## Abstract

Nutrient sensing and the subsequent metabolic responses are fundamental functions of animals, closely linked to diseases such as type 2 diabetes and various obesity-related diseases. *Drosophila melanogaster* has emerged as an excellent model for investigating metabolism and its associated disorders. In this study, we used live-cell imaging to demonstrate that the fly functional homolog of mammalian glucagon, Adipokinetic hormone (AKH), secreted from AKH hormone-producing cells (APCs) in the corpora cardiaca, stimulates intracellular Ca^2+^ waves in the larval fat body/adipose tissue to promote lipid metabolism. Further, we show that specific dietary amino acids activate the APCs, leading to increased intracellular Ca^2+^ and subsequent AKH secretion. Finally, a comparison of Ca^2+^ dynamics in larval and adult fat bodies revealed different mechanisms of regulation, highlighting the interplay of pulses of AKH secretion, extracellular diffusion of the hormone, and intercellular communication through gap junctions. Our study underscores the suitability of *Drosophila* as a powerful model for exploring real-time nutrient sensing and inter-organ communication dynamics.

## Introduction

Nutrient sensing and subsequent metabolic regulation are vital for the survival and well-being of any living organism. Among the multiple metabolic signals modulated by diverse nutrients, the evolutionarily conserved insulin-glucagon nexus plays a central role in metabolic homeostasis. The antagonistic role of insulin and glucagon is well established and the balance between them is precisely coordinated to maintain physiologically normal blood sugar levels. Dysregulation of this balance can result in debilitating metabolic diseases ranging from diabetes to glucagonomas.

Extensive studies have elucidated the molecular mechanisms regulating insulin and glucagon levels in response to dietary sugar and carbohydrate-mediated hormonal control. While much is known regarding the role of dietary sugars, hormonal regulation by dietary amino acids has received less attention, despite its potential significance in diabetes and cardiovascular diseases^1^. Increasing evidence suggests that dietary amino acids play a direct or indirect role in regulating insulin secretion by mammalian pancreatic islets^2–5^ Similarly, dietary amino acid-induced hyperglucagonemia in humans and other mammals has also been reported^6–11^. Further, studies in mammals indicate that a high protein diet triggers glucagon release from pancreatic alpha islet cells^12–15^. In addition, recent studies have revealed that several amino acids stimulate the secretion of glucagon from pancreatic α-cells which, in turn, promotes amino acid metabolism, gluconeogenesis, and lipid β-oxidation in the liver^16^. This regulation, referred to as the Liver-α-Cell axis, is considered to play important roles in metabolic diseases, including diabetes, obesity, and nonalcoholic fatty liver disease (NAFLD)^17^. Despite these studies, it remains to be determined whether amino acid sensing is evolutionarily conserved beyond mammals. Answering this question would provide valuable insights into the fundamental mechanisms of metabolism and its dysregulation in diseases, and, importantly, would enable studies of this underexplored process in model organisms.

In recent years, *Drosophila melanogaster* has emerged as a highly relevant model organism for investigating the influence of macro-nutrients, such as sugars, amino acids, and lipids, on insulin- and glucagon-dependent metabolic pathways^18–20^. Insects produce insulin-like peptides (DILPs) from the insulin-producing cells (IPCs) located in the brain, while Adipokinetic hormone (AKH), a functional homolog of glucagon, is synthesized and released from AKH-producing cells (APCs) located in the corpora cardiaca (CC). AKH plays a pivotal role in maintaining blood sugar homeostasis and modulating starvation-induced hyperactivity in adult flies^21,22^; however, its regulation remains less understood compared to that of insulin^23^. Various dietary, hormonal and neuronal factors regulating insulin and glucagon signaling have been identified in the fly^24^. Notably, it has been recently demonstrated that IPCs directly sense the dietary amino acid leucine, which consequently regulates the secretion of DILPs^25^. In contrast, whether specific amino acids exert regulatory control over AKH production and secretion has remained unknown.

In addition to the nutrient-dependent hormone release and hormonal control of peripheral metabolism, the temporal dynamics of nutrient sensing and inter-organ communication of metabolic hormones are inadequately understood. This includes the precise temporal kinetics of hormone release under different nutrient conditions and the response speed of the target organs to the hormone. In this study, we used live-cell imaging to study the real-time dynamics of nutrient-dependent hormone secretion and the response of adipose tissues in both *ex vivo* organ cultures and *in vivo* free-behaving animals. Using organ co-culture and organ-conditioned media, we identified AKH as the primary neuropeptide that physiologically activates Ca^2+^ waves in the larval fat body, a functional homolog of the mammalian liver and adipose tissue. Ca^2+^ waves are transient elevations of cytosolic Ca^2+^ that propagate between neighboring cells for signal transmission^26^. Previous studies have demonstrated that AKH induces Ca^2+^ increase in isolated insect fat body^27^; however, whether it is the primary hormone responsible for *in vivo* Ca^2+^ waves has remained unclear. In this study, we show that fat body Ca^2+^ waves are completely abrogated in *Akh receptor* (*AkhR*) mutant flies, suggesting that AKH is the primary hormone that triggers these Ca^2+^ waves *in vivo*. The clear relationship between AKH release and fat body Ca^2+^ waves provides a way to monitor the real-time release of AKH *in vivo*. Intriguingly, for the first time, we observed pulsatile secretion of AKH from the APCs, which initiates global Ca^2+^ waves spreading from the larva head to tail.

This observation is reminiscent of glucagon-induced Ca^2+^ waves observed in the human liver^28^. However, the biological implications of such wave-like Ca^2+^ activity within the liver remain elusive. To explore the significance of intercellular Ca^2+^ propagation, we genetically disrupted the gap junctions in the fat body, which effectively abolished intercellular Ca^2+^ transmission. To our surprise, interrupting the intercellular propagation of the Ca^2+^ signal led to contrasting changes in cytosolic Ca^2+^ at different developmental stages: Ca^2+^ levels were increased in the larval fat body but reduced in the adult fat body following the disruption of gap junctions. These changes correspondingly resulted in an increase in triacylglycerol (TAG) catabolism during larval stages and a decrease in adults. We hypothesize that the global Ca^2+^ waves present during the larval stage underlies these differences in dependence on intercellular Ca^2+^ diffusion. Finally, we found that secretion of AKH from the APCs is not only repressed by sugar, but also promptly stimulated within minutes after consumption of particular amino acids, and that consumption of these amino acids promotes fat loss in both larvae and adult flies through AKH-dependent signaling. Altogether, our study reveals that the control of glucagon/AKH signaling by specific amino acids is evolutionarily conserved, which in turn regulates metabolism in the fat body via elevating intracellular Ca^2+^ and its subsequent molecular targets.

## Materials and Methods

### Fly husbandry

Flies were raised on standard food (2000 ml water, 12.8 g agar, 80 g yeast, 112 g cornmeal, 176 g glucose, 2.5 g methylparaben, 20 ml propionic acid, total 2 l of fly food) at 25 °C with 12 h:12 h light: dark cycles. Fly strains are described in **Supplementary Table 1.**

### Ex vivo GCaMP imaging in APCs

Early 3^rd^ instar larval brains were dissected in the HL6 buffer (108 mM NaCl, 8.2LmM MgCl_2_, 4LmM NaHCO_3_, 1LmM NaH_2_PO_4_, 5 mM KCl, 2LmM CaCl_2_, 80 mM sucrose, 5 mM HEPES, pH 7.3) and were immobilized using a tissue holder (Warner Instruments) in the perfusion chamber. The samples in HL6 buffer were recorded for 1 min to generate a baseline. Next, the solutions were changed to HL6 bufferL+LAA (5 mM) with the pH adjusted back to 7.3 by gentle perfusion for 10 min. Solutions in the perfusion chamber were controlled by a valve commander (Scientific Instruments). After stimulation, samples were washed out again with HL6. All imaging studies were performed with a Leica M205 FCA high-resolution stereo fluorescence microscopy (Leica). Image analyses were performed in ImageJ and plotted in Excel (Microsoft).

### Ex vivo imaging of larval fat bodies

The *ex vivo* imaging chambers were assembled with a live cell imaging dish (Nest, 801001), a metal ring with an inner diameter of 10 mm, and a cellulose film from a tea bag modified to match the chamber. Briefly, the fat bodies of 3^rd^ instar larvae were dissected in Schneider’s medium (Sigma) and placed at the center of the imaging dish in 20 μl of Schneider’s medium. Next, the modified film was gently placed on top of the fat body and the metal ring gently placed on the film to trap the tissue under the film. Finally, 200 μl of Schneider’s medium was added inside the insert. The samples in Schneider’s medium were recorded for 5 min (time interval:5s) to generate a baseline. Then, the Schneider’s medium was replaced by Schneider’s medium containing 100 ng/mL of different synthesized *Drosophila* neuropeptides (Genscript, the sequences of the peptides are shown in **Supplementary Table 2**), and recording was done for 20 min (time interval:5s). Time-lapse recording was performed on a Leica DMi 8 equipped with a Leica DFC9000 sCMOS camera and a 1.25x HC PL Fluotar objective (Leica). Specially, for higher resolution, a 5x N Plan objective (Leica) was used in the gap junction knockdown experiments. GCamp5G was excited with a 475 laser. Imaging was performed in a dark room at 18 °C. All image analyses were conducted using ImageJ and MATLAB. We used MATLAB to remove background noise and preserve the edges of the sample.

### Ex vivo imaging of adult fat bodies

For adult flies, as most of the fat bodies adhere to the inside of the abdominal cavity, we dissected the dorsal shell together with the fat bodies. The dorsal shell was then adhered to a live cell imaging dish by Vaseline, so the fat bodies face upwards, bathed in 200 uL Schneider’s medium. Time-lapse recording was performed on a Leica M205 FCA high-resolution stereo fluorescence microscope (Leica) equipped with a Leica DFC7000 GT camera. The remaining steps and parameters are similar to those in the larval experiment.

### In vivo imaging of immobilized larvae and adults

For larvae, we attached two coverslips to a microscope slide with double-sided tape to form a thin slit, then used plasticine to plug both sides of the slit so that the gap in the slit is the same width as a 3^rd^ instar larva. Next, a coverslip was added on the top of the chamber to immobilize the larva. For adults, the six legs of an adult female fly were cut off, and the wings adhered to a live cell imaging dish with Vaseline so that the abdomen faced upward. Time-lapse recording was performed on a Leica M205 FCA high-resolution stereo fluorescence microscope (Leica) equipped with a Leica DFC7000 GT camera. All image analyses were performed using ImageJ and MATLAB.

### In vivo imaging of fat bodies of freely behaving larvae

We used a 3D printed mold (Wenext) to produce an agarose gel containing different nutrients. The center of the gel has a 11mm*13mm*0.66mm chamber, which can provide a free-behaving arena for more than 10 early 3rd instar larvae. Finally, a coverslip was added on top of the chamber to prevent the larvae from escaping. After being allowed to acclimate for 10 min, larvae were recorded for 20 min (time interval:5s), then all larvae were transferred to an agarose gel chamber containing another type of food for 20 min recording. In order to ensure that there was no food residue in and on the body of the 3^rd^ instar larvae, larvae were starved for 9 hours before imaging and washed with ddH20 during each transfer process. Time-lapse recording was performed on a Leica M205 FCA high-resolution stereo fluorescence microscope (Leica) equipped with a Leica DFC7000 GT camera. All image analyses were conducted using ImageJ and MATLAB. In image processing, we used connected component analysis to approximate each connectome as a larva. Fluorescent signal changes were normalized using the following formula: ΔF/F0L=L(F(t) – F0)/F0, where F(t) is the fluorescence at time t, F0 is the average baseline (before transfer). Plotting of graphs and statistical analyses were conducted with GraphPad Prism 8.4.2.

### In vivo imaging of APCs of freely behaving larvae

To perform high-resolution neuronal imaging of freely behaving 1^st^ instar larvae, we used an extended-depth-of-field microscope with two modules (built by Quan Wen): 1. A darkfield imaging module equipped with a 4X NA 0.2 air objective (Nikon, Japan) and a high-speed near-infrared camera (acA2000-340kmNIR, Basler ace), used to track and record a freely behaving 1^st^ instar larva. 2. A fluorescence imaging module equipped with a 10X NA 0.3 air objective and a sCMOS camera (Zyla 4.2, Andor Inc., UK), with an imaging surface split in two by Optosplit II (Cairn, UK), which allows simultaneous recording of two fluorescent signals (calcium-sensitive GCaMP and calcium-insensitive RFP used as reference). Through extended-depth-of-field technology, the effective depth of field was extended by about 5 times, avoiding errors on the Z-axis caused by motion. Before imaging, the 1^st^ instar larvae were starved for 6 hours, and then the larvae were gently picked with a brush into an agarose gel chamber (Φ20 mm*0.15 mm) made with a 3D printed mold (Wenext) for 15-min recording. All image analyses were conducted using ImageJ and MATLAB. The fluorescent signal changes were normalized using the following formula: ΔF/F0L=L(F(t) – F0)/F0, where F(t) is the fluorescence at time t, F0 is the average baseline (before transfer). Plotting of graphs and statistical analyses were conducted with GraphPad Prism 8.4.2.

### AKH secretion assay

To measure AKH secretion and retention, early 3^rd^ instar larvae were picked out from standard food and washed with ddH20, and after feeding for a period of time under different dietary conditions, the brains were dissected in PBS (1.86LmM NaH2PO4, 8.41LmM Na2HPO4, 175 mM NaCl) and fixed in 4% v/v paraformaldehyde (PFA)/PBS for 30 min at 23 °C. After washing in PBST (PBS + 0.05% Triton X-100, BBI) (3 times, 10 min each), the brains were blocked in 5% BSA (Solarbio, A8020) in PBST for 1 h at 25 °C, then incubated with rabbit anti-AKH (1:1000; ABclonal, WG-05853) for 12-20 h at 4 °C. After washing in PBST (3 times, 10 min each), the sample brains were incubated with Alexa Fluor 488 goat anti-rabbit IgG (1:500; Invitrogen) for 1 h at 25°C and washed again using PBST (3 times, 10 min each). We used an antifade agent to mount the samples. All images were acquired using a Leica M205 FCA high-resolution stereo fluorescence microscope (Leica). Quantifications analyses were conducted using ImageJ and MATLAB. Plotting of graphs and statistical analyses were conducted with GraphPad Prism 8.4.2.

### Wave direction analysis

We set the direction of the larvae from head to tail as columns, took the average of each row, and derived a one-dimensional vector for each frame. Next, we correlated these measurements with time to obtain a two-dimensional graph to show the direction of calcium wave movement. Image analyses were conducted using MATLAB.

### Ca^2+^ proportion calculation

We use the baseline fluorescence of GCaMP5G as the threshold to calculate the proportion of the area where calcium activation occurs. Each calculation requires at least 20 minutes of time-lapse data. This parameter also represents the probability of a calcium activation event occurring per unit time per unit area. Image analyses were conducted using MATLAB. Plotting of graphs and statistical analyses were conducted with GraphPad Prism 8.4.2.

### Wave velocity calculation

We first defined two lines parallel to the wavefront. Then, we measured the time required by the wavefront to cross the distance between the two lines to manually calculate wave velocity. Image analyses were conducted using ImgaeJ. Plotting of graphs and statistical analyses were conducted with GraphPad Prism 8.4.2.

### Ca^2+^ propagation calculation

To characterize calcium wave propagation, we artificially set a parameter “Ca^2+^ propagation area/s”. According to the concept of connected component in image processing, we calculated the area of each connected region where calcium activation occurred in the 20-min time-lapse data, and calculated the average value. Image analyses were conducted using MATLAB. Plotting of graphs and statistical analyses were conducted with GraphPad Prism 8.4.2.

### TAG assay

5 flies from each group were homogenized with 100 μl of isopropyl alcohol (BBI, A600918-0500), centrifuged at 10,000g for 10 minutes at 4°C, and the supernatant was collected. 2 μl of sample solution was mixed with 200 μl of assay reagent (Elabscience, E-BC-K261-M), and incubated at 37°C for 10 minutes. We measured the absorbance at 492 nm in a microplate reader (Thermo Scientific Multiskan FC). Plotting of graphs and statistical analyses were conducted with GraphPad Prism 8.4.2.

### Statistics

Sample sizes were determined through preliminary experiments and previous studies to ensure that the final results are statistically significant. All assays were repeated more than three times. Quantitative and statistical parameters, including statistical methods, error bars, n numbers, and p-values, are indicated in the legend of each figure. Error bars shown in all results are from biological replicates. Differences were assessed using a two-tailed unpaired Student’s t test. P < 0.05 was considered statistically significant. Significance was noted as *p < 0.05, **p < 0.01, ***p < 0.001.

## Results

### Periodic Ca^2+^ waves in fly larval adipose tissues are induced by a brain-derived factor

Ca^2+^ waves are complex signaling phenomenon that have been documented across multiple biological systems and play roles in multiple processes such as nutrient and hormone sensing, secretion, wound response, smooth muscle contraction and mechanical sensing^29–34^. However, the *in vivo* triggers of these Ca^2+^ waves are usually not known or not well characterized. The fly fat body, which is the functional equivalent of the mammalian liver and adipose tissue, plays central roles in metabolic regulation and nutrient sensing^35^. Ca^2+^ levels in the fly fat body have been reported to be essential in lipolysis^36,37^, and *in vivo* Ca^2+^ activities in the larval fat body have been observed^38^. Similarly, Ca^2+^ waves have also been reported in the mammalian liver^33^. Despite these findings, Ca^2+^ waves have not been systematically investigated *in vivo* in the fly fat body, and the triggering factor and biological function of Ca^2+^ waves remain unexplored. To address these questions, we expressed the genetically encoded Ca^2+^ indicator *GCaMP5G* in the larval fat body using fat specific *Lpp-Gal4* and immobilized larvae within a glass channel for observation (**Supplementary Fig. 1A**). To our surprise, we noticed global Ca^2+^ waves that emanate in a periodic fashion from the larval head to tail (**Fig. 1A-B, Supplementary video 1**). To ensure that these waves were not an artefact due to immobilization of the larvae, we also examined Ca^2+^ activity in freely behaving 3^rd^ instar larva. Strikingly, we observed similar prominent global Ca^2+^ waves, indicating that these waves were not a consequence of immobilization (**Supplementary Fig. 1B-C).**

**Figure 1.**
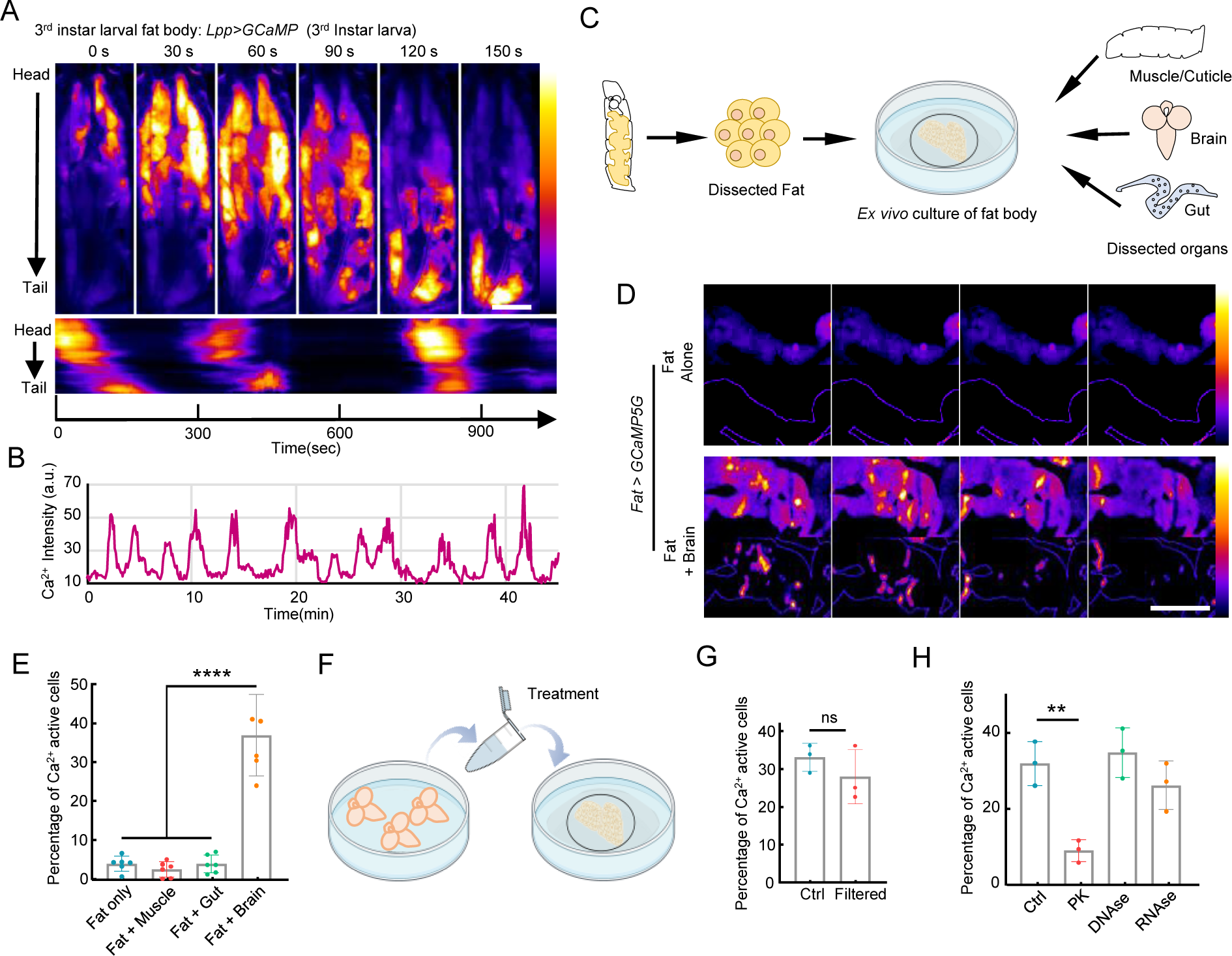
Periodic Ca^2+^ waves are triggered in the fly larval adipose tissues by a brain-derived factor. **(A-B)** A 3^rd^ instar larva was placed in a narrow glass channel to restrict its mobility. Adipose tissue specific Ca^2+^ activity was visualized using *Lpp-GAl4>UAS-GCaMP5G*. A kymograph of Ca^2+^ waves traveling from head to tail is shown. **(B)** Representative Ca^2+^ activity in a selected position of the larva. **(C)** Fat body was dissected and cultured in Schneider’s *Drosophila* medium with or without different larval organs. **(D)** Fat body cultured alone showed little Ca^2+^ activity, while fat body cultured together with dissected brains showed prominent Ca^2+^ waves. **(E)** Quantification of Ca^2+^ activities in cultured fat body under different conditions. **(F)** Brain-conditioned medium was used to treat the isolated fat body. **(G)** Brain-conditioned medium before and after filtration triggered significant Ca^2+^ waves in cultured fat bodies. **(H)** Ca^2+^ activity triggered by proteinase K-digested brain-conditioned medium was significantly reduced compared to untreated brain-conditioned medium. Proteinase K-digested brain-conditioned medium was treated with 0.1mg/mL proteinase K for 1 hour at 37°C and then filtered with Pierce 10K MWCO concentrator. The untreated brain-conditioned medium was processed similarly without addition of proteinase K. Data were plotted as mean ± SD. Scale bars, 500 μm (A), 500 μm (D).

Previous studies in fly imaginal discs have suggested that Ca^2+^ waves are triggered autonomously or in response to self-secreted epithelial-derived factors^30,39^. To discern whether the Ca^2+^ in the fat body is autonomously regulated or under the peripheral control of other organs, we dissected and cultured fat bodies in Schneider’s *Drosophila* medium. Interestingly, the isolated larval fat bodies displayed no Ca^2+^ waves except for a few cells that sustained damage during dissection and maintained an abnormally high Ca^2+^ level (**Fig. 1D**). These data suggest that fat body Ca^2+^ waves are regulated by other organs. To pinpoint the specific organ responsible for this regulation, we co-cultured fat bodies with different larval organs including muscle/cuticle, intestine, and brain (**Fig. 1C**). Interestingly, Ca^2+^ waves were only triggered when fat bodies were co-cultured with brains (**Fig 1. C-E, Supplementary video 2**), suggesting that these waves are initiated by a brain-derived factor.

To identify the causative factor, we generated a brain-conditioned medium by incubating Schneider’s *Drosophila* medium together with dissected larval brains (10 dissected 3^rd^ instar larval brains in 150 μl medium), and applied the conditioned medium to isolated fat bodies (**Fig. 1F**). As anticipated, the brain-conditioned medium induced robust Ca^2+^ waves (**Figure 1F-G**). To determine the nature of this factor, we filtered the brain-conditioned medium through a Pierce 10K MWCO concentrator that removed molecules larger than 10 kDa. This medium continued to evoke fat body Ca^2+^ waves, suggesting that the factor has a relatively small molecular weight (**Fig. 1G**). We then treated the conditioned medium with proteinase K (0.1mg/mL), DNAse I (1 U/mL) or RNAse A (0.1 μg /mL), followed by removal of these enzymes with the same 10 kDa filter. Among these treatments, only proteinase K significantly reduced the Ca^2+^ activities, suggesting that the factor is likely a small peptide (**Fig. 1H**).

### Fly Adipokinetic hormone (AKH) released from Akh-producing cells (APCs) is responsible for inducing Ca^2+^ waves in the fat body

To identify the relevant peptide(s), we screened 32 major fly peptides on isolated fat bodies (10 ng/mL each). Among them, only AKH evoked significant Ca^2+^ waves in the fat body (**Fig. 2A**). Removal of added AKH resulted in an immediate abrogation of Ca^2+^ activities, indicating that Ca^2+^ waves require a sustained supply of AKH (**Fig. 2B**). The observation that the fat body responds to the addition or removal of AKH within several seconds, together with the finding that there was no activity reduction in response to prolonged activation (up to 4 hours), suggests that the AKH receptor in the fat body has fast binding and dissociation kinetics without signal adaptation. Further, we found that the fat body is sensitive to AKH at doses as low as 0.1 ng/mL, which is within the physical range of neuropeptides. In addition, both the amplitude and frequency of Ca^2+^ oscillations generally increased with AKH concentration (**Fig. 2C, Supplementary Fig. 2A**). Finally, knocking down the only fly-encoded AKH receptor, *AkhR*, also reduced the Ca^2+^ oscillation frequency, suggesting that the frequency is regulated by the concentration of both ligand and receptor (**Supplementary Fig. 2B-C**).

**Figure 2.**
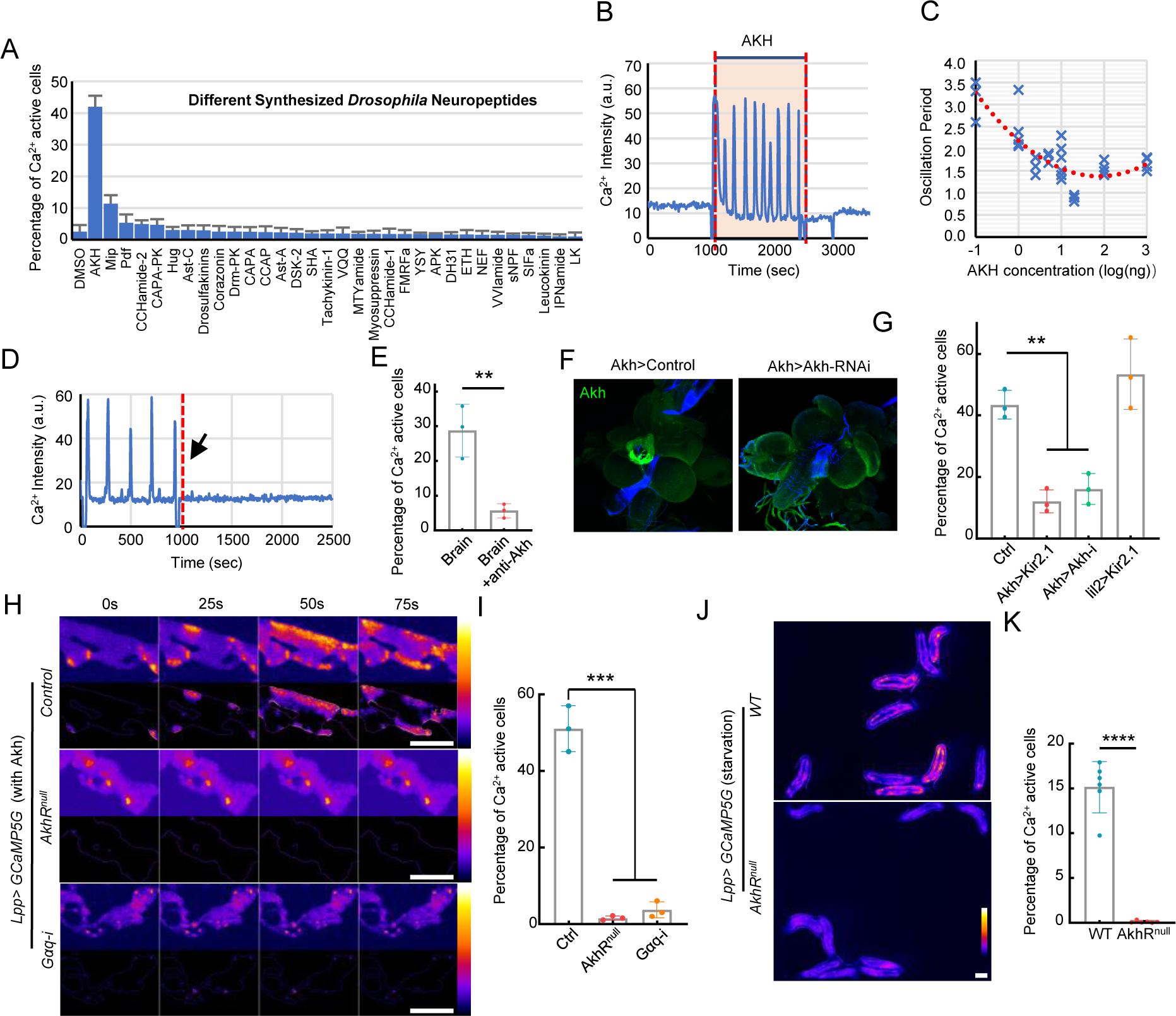
Brain-derived AKH is responsible for the Ca^2+^ waves in the fat body. **(A)** Ca^2+^ waves triggered by different synthesized fly neuropeptides (0.1 ug/mL) were quantified. **(B)** AKH (0.1 ug/mL) was added to the dissected fat body, incubated for 15 min, and then washed away. **(C)** Relationship between the period of Ca^2+^ oscillation and concentration of applied AKH. **(D-E)** Ca^2+^ activity triggered by brain-conditioned medium was blocked by addition of anti-AKH antibody (1:100 dilution). **(F-G)** Conditional medium derived from brains with AKH producing cells silenced by Kir2.1 or with *Akh* knockdown showed a significant reduced ability to trigger Ca^2+^ activity in the fat body. **(H-I)** Fat body with *AkhR* knockdown or *Gαq* knockdown failed to respond to AKH. **(J-K)** Ca^2+^ waves in free behaving larvae were significantly reduced in *AkhR* mutant larvae. Data were plotted as mean ± SD. Scale bars, 500 μm (H), 1000 μm (J).

To test whether AKH is indeed the fat stimulation factor in the brain-conditioned medium, we added an anti-AKH antibody in the medium to sequester free AKH. Ca^2+^ waves were blocked immediately after adding the anti-AKH antibody (**Fig. 2D-E**). Next, as AKH is secreted from Akh-producing cells (APCs) of the ring gland, which is associated with the larval brain, we knocked down *Akh* in the APCs (*Akh-Gal4>UAS-Akh-RNAi*) or inhibited AKH release by expressing the Kir2.1 potassium channel (*Akh-Gal4>UAS-Kir2.1)*. Brains unable to release AKH resulted in a significant decrease in triggering of fat body Ca^2+^ waves in the co-culture experiment (**Fig. 2F-G**). Finally, fat bodies from *AkhR* mutant larvae no longer responded to the applied AKH peptide (**Fig. 2H-I, Supplementary Video 3**) and the Ca^2+^ waves were completely blocked in *AkhR* mutant larvae *in vivo* (**Fig. 2J-K, Supplementary Video 4**).

AkhR is a GPCR receptor that has been found to trigger both cAMP and Ca^2+^ ^35^. Previous studies have shown that AKH regulates Ca^2+^ increase by activating phospholipase C (PLC) and subsequent generation of IP3 and DAG^27^. To block the AKH-induced Ca^2+^ increase, we knocked down *G*_α_*q,* which is responsible for the GPCR-dependent PLC activation. Knocking down *G*_α_*q* in fat bodies blocked the AKH-triggered Ca^2+^ waves as observed in *AkhR* mutant animals, suggesting that Gαq is a required AKH-downstream regulator for the Ca^2+^ activity (**Fig. 2H-I**). This is further supported by *G*_α_*q* overexpression, which generated active Ca^2+^ waves both in isolated fat bodies without the addition of AKH and in freely behaving larvae on a 5% m/v sucrose diet (**Supplementary Fig.3A-B**). Conversely, knocking down *SERCA*, an important ER-located Ca^2+^ pump, led to a sustained elevation of Ca^2+^ in the fat body (**Supplementary Fig.3C-D**). Collectively, these findings suggest that Ca^2+^ activity in the larval fat body is triggered by AKH secreted from the APCs, which depends on Gαq-mediated efflux of Ca^2+^ from the ER downstream of the AKH receptor.

### Regulation of AKH release from APCs by amino acids

Our findings demonstrate that the fat body exhibits a specific and rapid response to extracellular AKH. This property provides a clear and direct readout for monitoring real-time secretion of AKH in freely behaving larvae under different nutrient conditions. As most previous live-imaging setups used immobilized or even anesthetized larvae, which may cause undesirable distress and artefacts, we decided to study metabolic signaling in freely behaving animals. In addition, as previous studies have suggested that AKH is a functional homolog of mammalian glucagon, which is released upon starvation^22,40–42^, we used early 3rd instar larvae first starved for 9 hours to cleanse their digestive systems then fed on 2% sucrose for 20 min, which we found brings the fat body Ca^2+^ activities and AKH secretion to a basal level (**Fig. 3A**). Subsequently, these conditioned larvae were transferred onto 2% agarose plate with 2% sucrose, 2% sucrose plus 10% Tryptone (a protein-rich diet equivalent to approximately 5% protein), or 2% agarose with no nutrient (starvation diet). Larvae fed on a starvation diet displayed a significant increase in fat body Ca^2+^ waves after ∼10 minutes of feeding, indicating a swift response to starvation, which agrees with the established function of AKH as a starvation-induced hormone like mammalian glucagon. Importantly, consumption of the protein-rich diet also significantly increased Ca^2+^ activities in the fat body within just 10-15 minutes (**Fig. 3B-C, Supplementary Video 5**), suggesting that AKH secretion is also triggered by amino acids. To confirm whether the protein-induced fat body Ca^2+^ waves are indeed AKH-dependent, we compared WT larvae with larvae carrying a *AkhR* mutation or fat-body specific *AkhR* knockdown using *Lpp-Gal4*. In both cases, we observed a significant reduction of the Ca^2+^ waves under both starvation and protein-feeding conditions, supporting the idea that both processes depend on AKH signaling (**Fig. 3D-E**).

**Figure 3.**
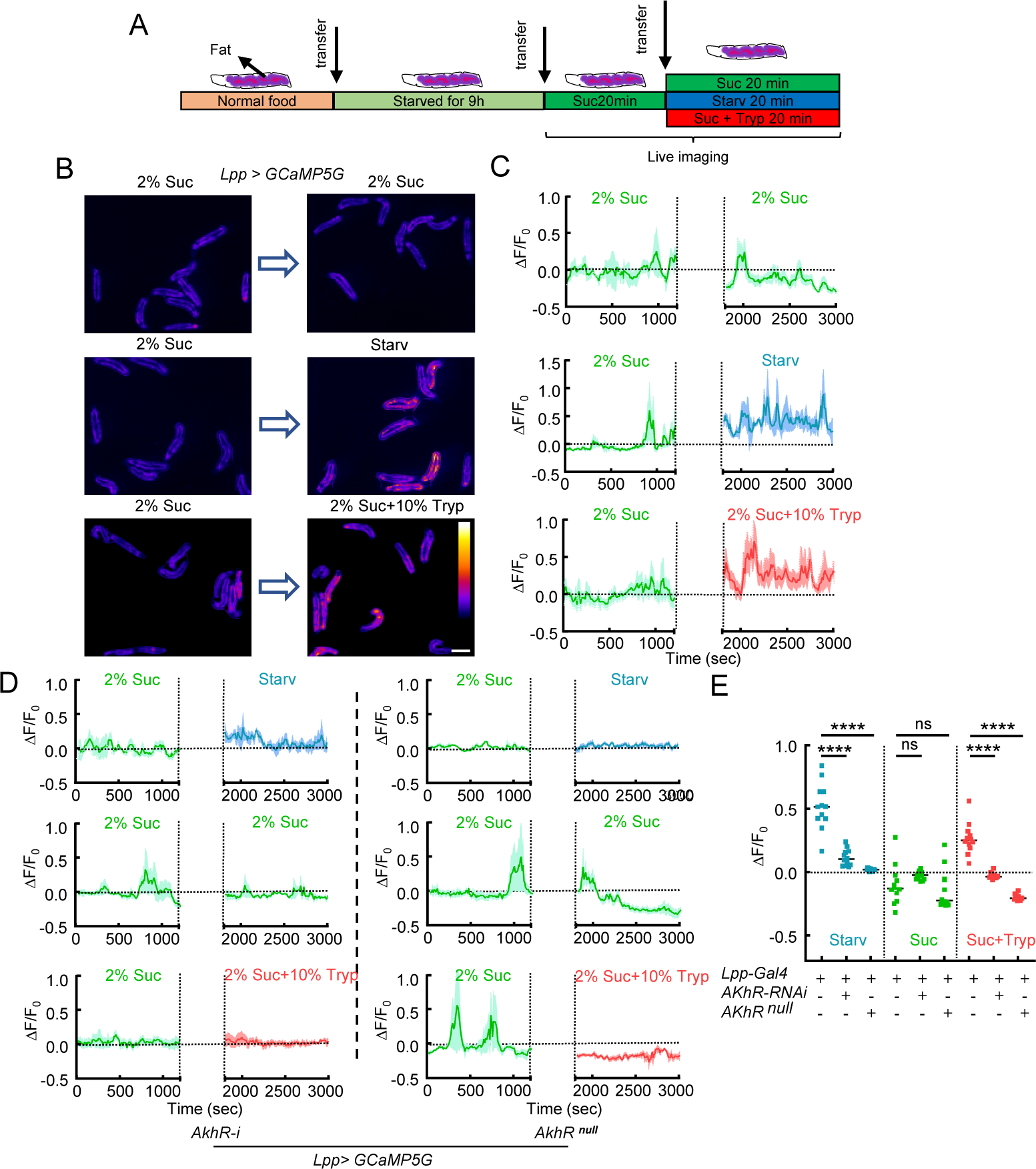
Fat AKH-dependent Ca^2+^ waves are regulated by dietary sugar and protein. **(A)** Early 3^rd^ instar fly larvae were transferred onto 2% agarose for 9 hours and then onto 2% Agarose + 2% sucrose for 20 min. These conditioned larvae were further transferred onto different test food containing 2% sucrose, 2% sucrose + 10% Tryptone (protein diet), or no nutrient (starvation diet). **(B-C)** Representative Ca^2+^ images and Ca^2+^ activities of free behaving larvae transferred between different diets. Ca^2+^ activities were calculated from the average signals obtained from all imaged larvae. The gap between two diets is the larvae transfer time which is ∼10 min. **(D)** Representative Ca^2+^ activities in the fat body of free behaving larvae with whole-body *AkhR* mutant or fat specific knockdown of *AkhR*. **(E)** Quantification of Ca^2+^ activities in the fat body of free-behaving fly larvae. Data were plotted as mean ± SD. Scale bars, 2000 μm (B).

The *in vivo* Ca^2+^ activity of the fat body suggests that AKH release from APCs is triggered by amino acids. To test this directly, we examined whether APC activity is regulated by a protein diet in freely behaving larvae. We used *Akh-Gal4>GCaMP5G-T2A-mRuby3* to visualize and quantify Ca^2+^ activity in the APCs. However, as the thick cuticle of the 3^rd^ instar larvae caused a strong blurring of the signal from the APCs when they moved, we decided to use 1^st^ instar larvae, which are much smaller and more transparent. Although imaging was improved, imaging the APCs in freely behaving 1^st^ instar larvae was still challenging. To improve both image resolution and imaging speed, we used an Extended-Depth-of-Field (EDoF) microscope, which turns the slow 3D imaging into a quick 2D acquisition (**Supplementary Fig.4A-D**). In addition, because the Ca^2+^ signals from the APCs are much dimmer than those of the fat body, we used a starvation diet as the initial condition to achieve a reliable visualization of Ca^2+^ activity. 1^st^ instar larvae were starved for 6 hours on 2% agarose plate, then transferred to new agarose plates containing 5% sucrose (sugar diet), 10% Tryptone (protein-rich diet), or 2% agarose only (starvation) (**Fig. 4A**). As expected, Ca^2+^ levels in the APCs significantly decreased after being fed on 5% sucrose (**Fig. 4B-G, Supplementary Video 6**). Interestingly, Ca^2+^ levels in the IPCs were reduced when we performed simultaneous dual labeling of both the APCs and IPCs (**Supplementary Fig.4E-F**). This real-time observation of the counter activities of the IPCs and APCs strongly supports the reliability of this live-imaging system. Next, we tested the effect of a 10% tryptone diet and found that protein consumption triggers a Ca^2+^ increase in the APCs after ∼10 minutes of feeding, which is consistent with the quick Ca^2+^ response observed in the fat body (**Fig. 4B-G**). Note that we used 5% sucrose instead of 2% because 5% sucrose triggers a quicker and stronger AKH suppression. However, for the experiment in 3^rd^ instar larvae, the secretion of AKH secretion is too severely suppressed with 5% sucrose, making the Tryptone diet less effective.

**Figure 4.**
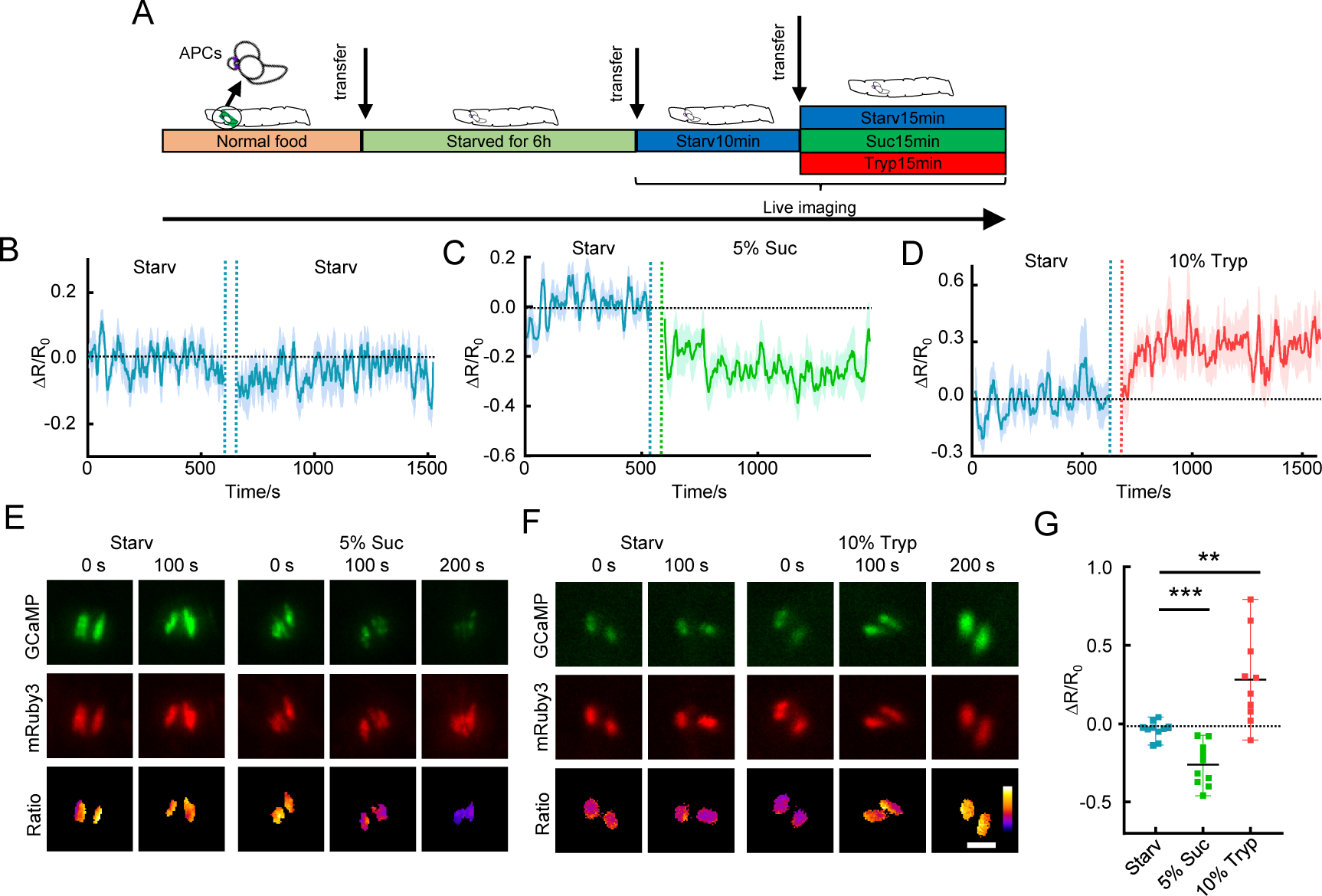
AKH secretion from APCs is regulated by amino acids. **(A)** 1^st^ instar fly larvae were transferred onto 2% agarose for 6 hours and then onto different test food containing 5% sucrose, 10% Tryptone (protein diet), or no nutrient (starvation diet). **(B-D)** Representative Ca^2+^ activities in the APCs of free-behaving larvae transferred between different diets. The gap between the two diets is the larvae transfer time which is ∼10 min. **(E-F)** Representative Ca^2+^ images of the APCs of free-behaving larvae. **(G)** Quantification of Ca^2+^ activities in the APCs of free behaving larvae. Data were plotted as mean ± SD. Scale bars, 25 μm (E, F).

Studies in mammals have found that only some amino acids trigger glucagon release, with branch-chained amino acids failing to induce such secretion^16^. Thus, we tested which particular amino acids may activate AKH-mediated Ca^2+^ waves in *ex-vivo* fat body tissues using our conditioned-medium system. As the complete Schneider’s *Drosophila* medium contains all amino acids and triggers the release of AKH from dissected larval brains (**Fig. 1F-H**), we prepared a basal *Drosophila* HL6 buffer devoid of any amino acids (referred to as HL6(AA-) buffer). Subsequently, each amino acid (5 mM) was individually added to the HL6(AA-) buffer to assess its capacity to induce AKH release (**Fig. 5A**). Brain-conditioned HL6(AA-) buffer does not activate the cultured fat body; however, the addition of most small polar amino acids triggered AKH secretion and subsequent elevation of fat body Ca^2+^ **( Fig. 5B-C)**. Notably, the large branch-chained amino acids leucine (Leu) and isoleucine (Ile) failed to trigger AKH release, akin to their effects in the mammalian glucagon system. We further monitored Ca^2+^ activities in APCs in isolated larval brains. APCs respond to threonine (Thr) within two minutes and reach an activation plateau at around six minutes, consistent with the response speed observed *in vivo*. In contrast, APCs showed little response to Leu (**Fig. 5E-G**). Finally, the release of AKH from APCs was confirmed by AKH staining of APCs in 3^rd^ instar larvae. Starvation, known to trigger AKH release, served as a positive control (**Supplementary Fig. 5**). Next, we tested 36 hours of feeding on 5% sucrose, 5% sucrose plus 40 mM methionine (Met), or 5% sucrose plus 40 mM Leu (**Fig. 5H-J**). Met feeding significantly reduced the AKH signal in the APCs compared to the sugar control and Leu, suggesting that Met triggers the release of AKH *in vivo.* Previous studies have shown that increased AKH levels lead to lipolysis in fat body tissues, especially in adult flies^41^. Hence, we examined whether amino acid feeding could reduce triacylglycerol (TAG) content. Indeed, 40 mM Met feeding reduced the TAG content in adult flies, and this reduction was entirely abrogated in *AkhR* mutant animals (**Fig. 5K**). Altogether, our results show that specific dietary amino acids, i.e., Met and Thr, are sensed by APCs to trigger AKH release, which in turn activates Ca^2+^ waves in the fat body.

**Figure 5.**
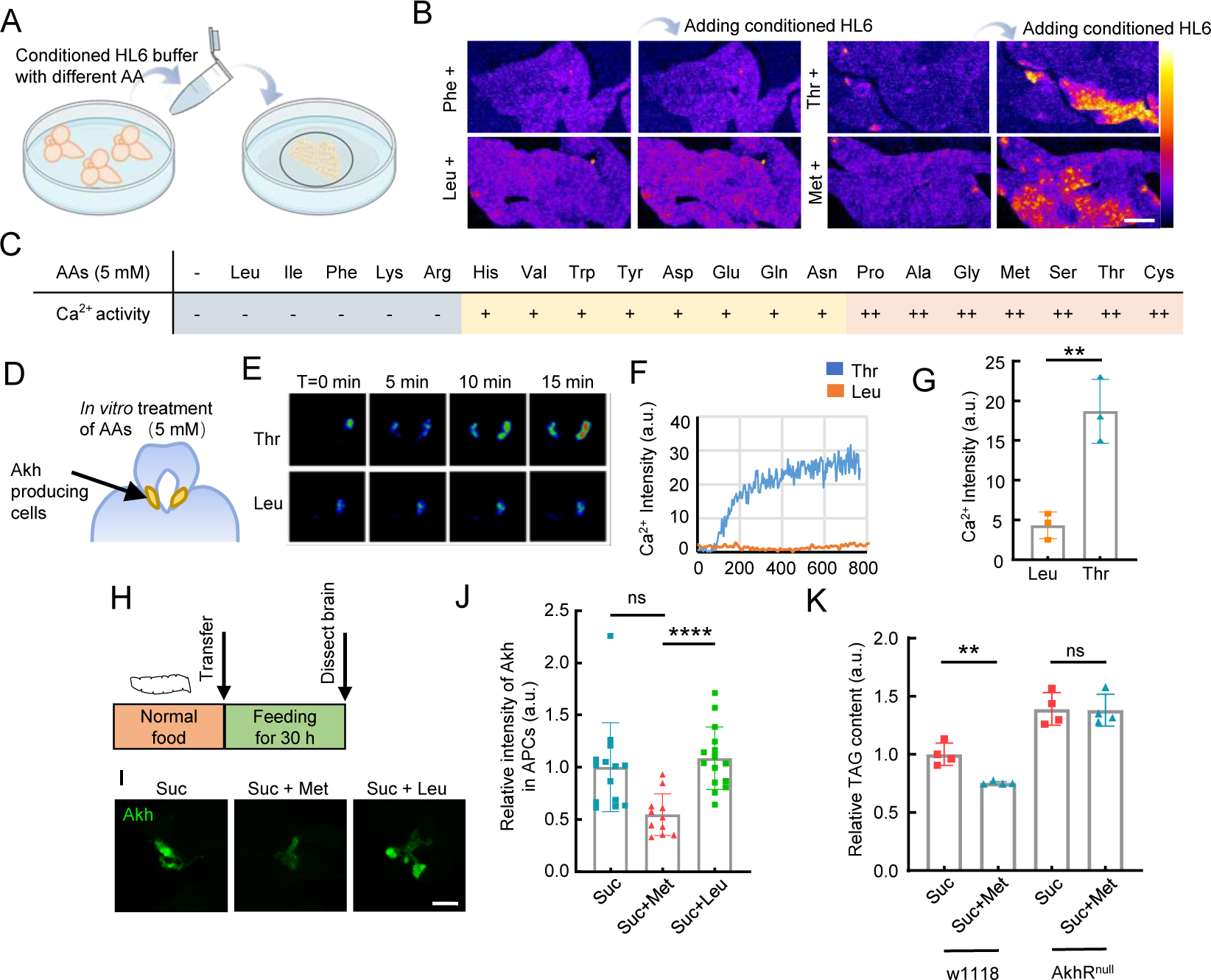
AKH secretion from APCs is regulated by amino acids. **(A)** Brain-conditioned HL6 media with each individual amino acid (5 mM) were generated and transferred to the dissected 3rd instar larval fat body. **(B)** Representative Ca2+ images of fat bodies before and after addition of brain-conditioned HL6 media with indicated amino acids. **(C)** Ca2+ activities in the bodies treated by brain-conditioned HL6 medium with indicated amino acids. Percentage of fat body cells with positive Ca2+ signal was indicated: “++” means more than 10%, “+” means 10%-5%, “-” means less than 5%. **(D)** AKH producing CC cells located at the “foot” region of the ring gland associated with the larval brain. Ca2+ activity in the CC cells were monitored using *Akh-Gal4>GCaMP5G*. **(E-F)** Dissected larval brains together with ring glands were incubated in HL6 medium with amino acids. Representative Ca2+ activities in APCs after addition of the indicated amino acids (5 mM) are shown. **(G)** Quantification of Ca2+ activities in the APCs 15 mins after the addition of the indicated amino acids. **(H)** 3rd instar larvae fed on indicated food for 36 hours and then dissected. AKH levels in the APCs were stained with anti-AKH antibody. **(I-J)** Representative stainings of AKH and quantifications of the fluorescent intensity in APCs of fly larvae fed on 2% sucrose, 2% sucrose + 40 mM Met, or 2% sucrose + 40 mM Leu. **(K)** TAG contents of adult flies fed on indicated foods for 48 hours were measured. Diets containing 2% sucrose or 2% sucrose + 40 mM Met were used. Data were plotted as mean ± SD. Scale bars, 50 μm (B), 25 μm (E, I).

### Global and local Ca^2+^ waves are regulated by distinct mechanisms

The AKH-induced Ca^2+^ waves observed in the fly fat body resemble the glucagon-triggered Ca^2+^ waves observed in the mammalian liver^43^. However, the significance of the wave-like nature of Ca^2+^ activities in both liver and fat body has not been explored. Meanwhile, the AKH-induced global Ca^2+^ waves propagating from the larval head to tail represent a previously undocumented phenomenon, with the underlying molecular mechanism uncharacterized (**Fig. 6A**). To investigate the biological implication of the intercellular Ca^2+^ propagations, we genetically blocked the diffusion of Ca^2+^ in the fat body. As most previous studies suggested that the intercellular Ca^2+^ transmission are mediated by gap junctions^26^, we screened all major gap junction proteins by RNAi and found that only *Inx2* and *Inx3* knockdown significantly scattered the intercellular Ca^2+^ waves in *ex vivo* fat bodies (**Fig. 6A-B, Supplementary Fig. 6**). This observation is consistent with previous studies that Inx2 is predominately required for gap junction function in the fly epithelium^26,30^, and that Inx3 probably functions together with Inx2 to form a heterohexamer^44^. Since *Inx2* knockdown exhibited the strongest phenotype, we used *Inx2-RNAi* to disrupt gap junctions in all subsequent experiments.

**Figure 6.**
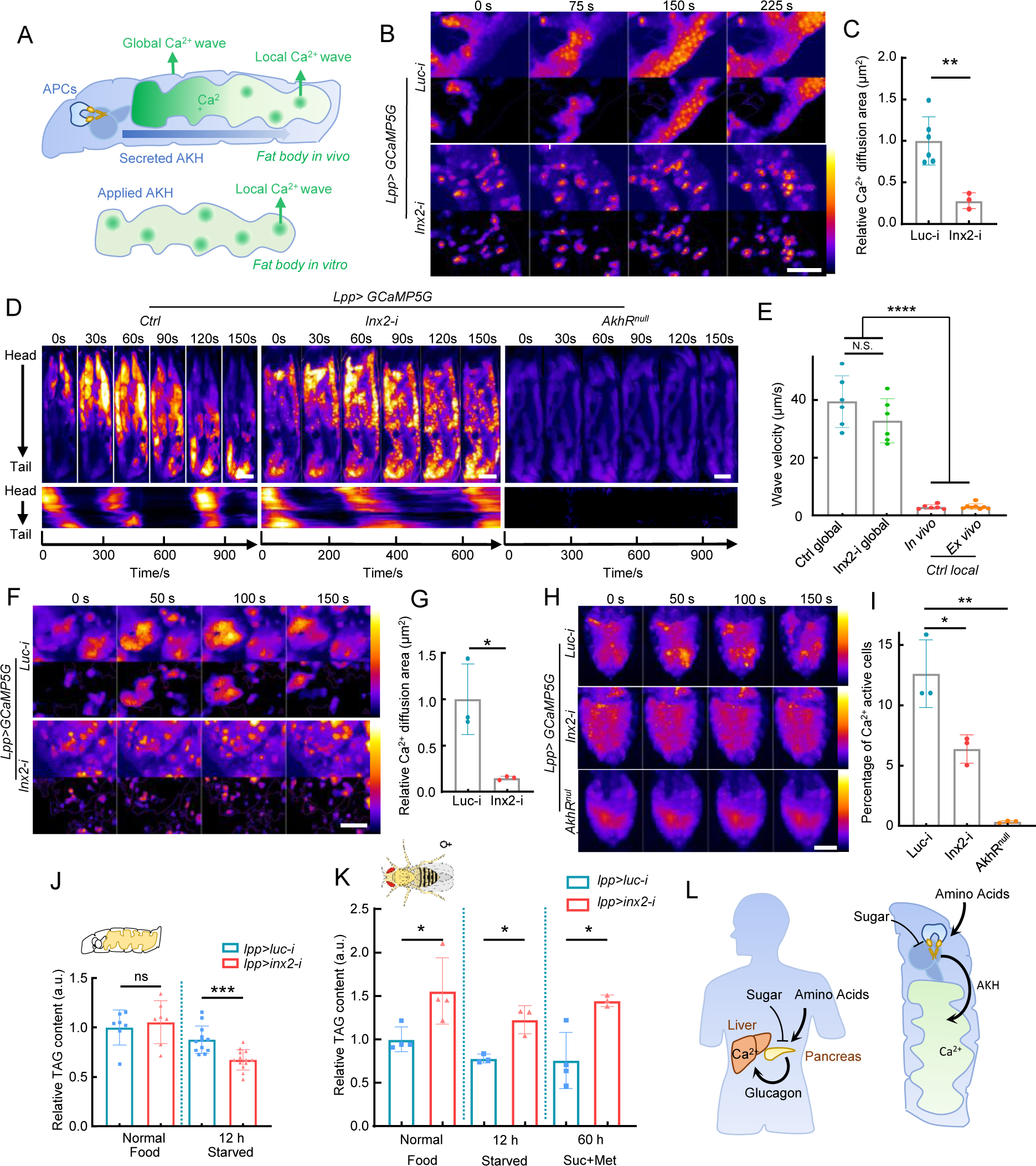
Global and local Ca^2+^ waves are generated by different mechanisms. **(A)** Both global and local Ca^2+^ waves are present in the larval fat body. The tissue-level Ca^2+^ waves traveling from larval head to tail is only observed *in vivo*. Conversely, local Ca^2+^ waves, initiated randomly in the fat body cells, are evident in both *in vitro* and *in vivo* experimental settings. **(B-C)** Disruption of intercellular gap junctions by *Inx2* knockdown significantly scattered the Ca^2+^ activities in *ex vivo* cultured fat body. **(D-E)** *Inx2* knockdown does not affect the magnitude or period of global Ca^2+^ waves *in vivo*, while in *AkhR* mutant both global and local Ca^2+^ waves are completely blocked. Larvae were kept under starved condition to trigger the release of endogenous AKH. **(E)** Quantification of the travelling velocity of the global and local Ca^2+^ wave. **(F-G)** *Inx2* knockdown completely scattered the Ca^2+^ waves in dissected adult fly fat bodies. **(H-I)** *Inx2* knockdown significantly reduced the Ca^2+^ waves in the adult fat body *in vivo*. **(J-K)** Effects of gap junction disruption in fat bodies during the larval and adult stages on TAG metabolism. **(L)** Sugar and amino acid sensing properties of AKH as well as its downstream Ca^2+^ waves are functionally conserved compared with mammalian glucagon. Data were plotted as mean ± SD. Scale bars, 500 μm (A), 500 μm (C), 200μm (F), 500 μm (H).

*In vitro* experiments demonstrated that *Inx2* knockdown effectively blocked the local propagation of Ca^2+^ waves (**Fig. 6B-C**). However, the *in vivo* global Ca^2+^ waves were surprisingly unaffected by *Inx2* knockdown (**Fig. 6C-D, Supplementary Fig. 7A**). In fact, Ca^2+^ oscillations in *Inx2-RNAi* flies exhibited an increase in amplitude and a decrease in frequency (**Supplementary Fig. 7B-D**). Similarly, the overall effect of *Inx2* knockdown led to an unexpected elevation of overall Ca^2+^ activity in free-behaving larvae (**Supplementary Fig. 7E-G**). To rule out the possibility of non-specific effects of *Inx2-RNAi*, we also tested the effects of the gap junction inhibitor carbenoxolone. Consistent with *Inx2-RNAi,* carbenoxolone treatment showed a similar increase of Ca^2+^ amplitude and frequency (**Supplementary Fig. 7H-J**).

Despite these changes, our data clearly demonstrate that the spreading of the global Ca^2+^ waves does not require intercellular Ca^2+^ diffusion. However, as both global and local Ca^2+^ waves were completely abrogated in *AkhR* mutants, suggesting that these Ca^2+^ activities in the fat body are triggered by extracellular AKH (**Fig. 6D-E**). Thus, we propose that local Ca^2+^ waves are generated by diffusion of Ca^2+^ from cells stochastically activated by AKH, but that the global Ca^2+^ waves are generated in response to the pulsed secretion and diffusion of a high level of extracellular AKH from the APCs near the larval head region (**Fig. 6A**). Supporting this hypothesis, we found that global Ca^2+^ waves travel approximately 10 times faster than local Ca^2+^ waves (**Fig. 6E**). These measurements indicate that the diffusion speed of secreted AKH in the larval hemolymph is around 40 μm/sec, which aligns with the free diffusion speed of a 1 kDa molecule (the size of AKH) in water.

Besides the larval fat body tissue, we also examined Ca^2+^ activity in the adult fat body. *Inx2* knockdown also completely blocked the intercellular diffusion of Ca^2+^ in the adult fat body (**Fig. 6F-G**). Notably, contrary to the larval fat body, the overall Ca^2+^ activities in the adult fat body were significantly reduced in adult flies with *Inx2* knockdown (**Fig. 6H-I**). This discrepancy is further supported by the opposite lipolytic phenotypes of *Inx2* knockdown in larvae and adults. As increase of AKH secretion and cytosolic Ca^2+^ level has been linked with TAG accumulation in adipose tissues^35–37,41^, we assessed the effect of gap junction disruption on TAG catabolism. TAG levels in *Inx2* knockdown larvae remained unaffected under standard feeding conditions, consistent with the fact that AKH is not secreted under fed conditions (**Fig. 6J**). TAG levels in gap junction mutant larvae were significantly lower than in control larvae following starvation (**Fig. 6K**), which aligns with our observation that gap junction blockage increases AKH-induced Ca^2+^ activities in larvae. In contrast, adult flies with disrupted gap junctions in the fat bodies displayed a significant increase in TAG accumulation under normal diet, starvation, and amino acid feeding conditions (**Fig. 6L**), consistent with the reduction of Ca^2+^ activities in the adult fat body. We speculate that the strikingly different responses of larval and adult fat bodies to gap junction disruption could be attributed to the presence of global Ca^2+^ waves in larvae.

## Discussion

In this study, we demonstrate that AKH secreted from APCs stimulates intracellular Ca^2+^ waves in the *Drosophila* fat body to promote lipid metabolism. In addition, we found that specific dietary amino acids activate the APCs, leading to increased intracellular Ca^2+^ and subsequent AKH secretion. Furthermore, we discovered that the global and local Ca^2+^ waves in the fat body are generated through different molecular mechanisms: whereas the global Ca^2+^ wave is a result of extracellular diffusion of AKH, the local Ca^2+^ waves are formed through the intercellular diffusion of Ca^2+^.

### Ca^2+^ waves in the fly fat body are regulated by AKH-AkhR signaling

Previous studies have established Ca^2+^ as a key regulator of lipolysis in the fly fat body^36,37^, and AKH has been found to trigger Ca^2+^ increase in the fat body under *ex vivo* conditions^27^. Our current study demonstrates that the primary driver of Ca^2+^ activity is the AKH-AkhR pathway and its downstream effector *G*_α_*q*. Similar to mammalian glucagon, AKH is a central hormone that integrates diverse biological processes, including mobilization of lipid storage, stimulation of locomotion, oxidative stress protection, and immune response^35^. Our findings suggest that Ca^2+^ signaling within the fat body serves as a reliable real-time indicator for studying these processes in free-behaving animals.

### Difference in AKH-mediated Ca^2+^ propagation in the larvae and adult fat body

Ca^2+^ has been found to generate a variety of inter and intra-cellular activities, including flashes, sparkles, oscillations and waves, which have been implicated in diverse biological processes^26^. The global Ca^2+^ waves uncovered in our study present an intriguing model illustrating how an hormone can function as an extracellular orchestrator, creating a collectively moving pattern across a large epithelial tissue. Previous studies of Ca^2+^ waves have suggested that they rely on intercellular gap junctions to maintain functional Ca^2+^ levels^26,30,34^. However, our findings reveal that the global Ca^2+^ waves in the larval fat body bypass the need for intercellular Ca^2+^ propagation and are self-sufficient in maintaining tissue-level Ca^2+^ activities. These global Ca^2+^ waves also suggest that AKH is secreted in a strong pulsatile manner in larvae, a phenomenon similar to the pulsatile release of mammalian glucagon and insulin, which have been found to be disrupted in patients with type-2 diabetes and potentially contribute to hyperglucagonemia^45,46^. However, the biological significance of this pulsatile hormone release compared to continuous release is not clear. Our data suggest that in the fly larva, the pulsatile secretion of AKH may create a strong increase of hormone “shock” that collectively activates fat body cells, which may render the intercellular Ca^2+^ spreading between neighboring cells less important.

Intriguingly, the adult fat body depends on gap junctions to uphold a functional Ca^2+^ level under starvation or amino acid feeding conditions. Meanwhile, no organ-level global Ca^2+^ waves were observed—the Ca^2+^ waves detected *in vivo* in the adult fat body appear completely random, suggesting that AKH secretion may not be strong enough to collectively activate the fat body cells. Thus, in the absence of a strong extracellular AKH pulse, cells activated by AKH become sparsely distributed as in the adult fat body, requiring intercellular spreading of Ca^2+^. The biological significance behind these differences between larval and adult tissue remains an open question. Understanding these differences might help us understand the difference in response to AKH-AkhR signaling in larva vs. adult fly, i.e., AKH-AkhR signaling activation of lipolysis in adult fly fat body but not in the larval fat body.

### Dietary amino acid-mediated activation of APCs

The *in vitro* kinetics of Ca^2+^ activities also reveal intriguing aspects of amino acid sensing and AKH-AkhR signaling. We found that cultured APCs do not respond immediately to the applied amino acids but instead exhibit a gradual increase in activity over a 5-10 minute period. This suggests that amino acids may not function through a rapid response mechanism such as ligand-gated channels, but via a comparatively slower metabolic process, possibly involving an increase in cytosolic ATP following amino acid breakdown. However, it remains to be elucidated whether APCs sense amino acids directly or indirectly through other neurons or a secondary metabolite within the brain. Furthermore, the fast and continuous response of the fat body to extracellular AKH without adaptation implies that AkhR probably does not undergo activation-induced inactivation or endocytosis. This immediate and persistent response to AKH could provide flies with an advantage in promptly adapting their metabolic state to environmental changes.

By tracing Ca^2+^ activities in the fly fat body and AKH-producing cells in response to different nutrients, we found that AKH secretion is regulated by certain dietary amino acids in the fly hemolymph, which subsequently increases the mobilization of fat body lipids through AkhR-Gαq signaling. Interestingly, previous studies have identified other amino acids sensing mechanisms: amino acid triggers the release of GBP1/2 and Stunted from the fat body to stimulate the insulin-like peptides secretion and promote larval growth^47,48^, essential amino acids promotes the release of CNMa from intestine cells to regulate feeding behavior^49^, FMRFa secretion from brain neurons is triggered by amino acid consumption to mobilize lipid stores^50^, and insulin-producing cells (IPCs) sense amino acids to increase insulin-like peptides production^51^. It will be interesting to explore how and where these amino acid-dependent signals interact or integrate to achieve metabolic homeostasis.

Although we have observed that amino acids regulate the secretion of AKH, the precise biological significance of this phenomenon is still not fully understood. Our study primarily focused on the regulation of AKH secretion and its effect on fat body Ca^2+^ increase and subsequent TAG lipolysis. However, it is conceivable that the activation of AkhR has a more extensive role than facilitating neutral lipid mobilization. Recently, AKH has been found to activate extracellular signal-regulated kinase (ERK), which in turn increases amino acid catabolism and gluconeogenesis in the fly fat body^52^. Together with our observations, it seems plausible that amino acid-induced AKH secretion serves as a mechanism for flies to process excessive amino acids ingested from the diet. AKH is believed to be a functional homolog of mammalian glucagon, which is specifically produced under starved or low-energy conditions to promote lipolysis in peripheral organs. However, we found that AKH is not only stimulated under energy challenges conditions but also by a high protein diet. While a high protein diet has also been reported to trigger mammalian glucagon, which specific amino acids are involved is not known. A more comprehensive study of the downstream signaling pathway of AKH is needed to fully understand the consequence of this AKH-mediated amino acid sensing axis. In addition, our and previous work suggest that different dietary amino acids can trigger different neuronal centers in the fly brain i.e., leucine and isoleucine specifically trigger IPCs in the larval brain whereas we found that these two amino acids do not trigger APCs to release AKH. Further research is required to decipher how seemingly complex activation of different neuronal centers may take place following a high protein diet consisting of different groups of amino acids.

Lastly, the increase of amino acid uptake and usage by AKH signaling in the fat body resembles the recently discovered mammalian Liver-α-Cell axis whereby an increase in glucagon levels upregulates the expression of specific amino acid transporters such as Slc38a4 and Slc38a5 in the liver, thereby enhancing amino acid uptake and promoting gluconeogenesis as well as urea production^17^. It would be interesting to investigate whether certain amino acid transporters are similarly upregulated by AKH in the fly fat body. Moreover, elevated amino acid levels in the bloodstream not only stimulate glucagon secretion but also contribute to α-cell proliferation, leading to pancreatic α-cell hyperplasia in mice, creating a lasting endocrine feedback^17^. Whether the number or function of APCs are modulated by amino acid consumption in *Drosophila* is still unknown. Experiments examining whether high protein intake induces lasting effects on APCs could provide evidence for conservation of the Liver-α-cell axis across species.

## Supplementary Materials

**Supplementary Figure 1-7. Supplementary Video 1**: Ca^2+^ activities in immobilized 3^rd^ instar larva. **Supplementary Video 2**: Ca^2+^ activities in isolated larval fat body tissue cultured with or without the presence of brain tissues. **Supplementary Video 3**: Ca^2+^ activities in isolated fat body tissue from wild-type (WT), *AkhR* mutant, and *G*_α_*q* knockdown larvae. **Supplementary Video 4**: Ca^2+^ activities in the fat body of freely behaving WT and *AkhR* mutant larvae. **Supplementary Video 5**: Ca^2+^ activities in the fat body of freely behaving larvae as they are transferred from a 2% sucrose diet to indicated diets. **Supplementary Video 6**: Ca^2+^ activities in the APCs (AKH-producing cells) of 1^st^ instar larvae when transferred between indicated diets. **Supplementary Video 7**: Ca^2+^ activities in fat body treated with brain-conditioned HL6(AA-) medium supplemented with indicated amino acids. **Supplementary Video 8**: Ca^2+^ activities in isolated fat body from WT and *Inx2* fat body knockdown larvae. **Supplementary Video 9**: Ca^2+^ activities in immobilized 3^rd^ instar larvae expressing *Inx2-RNAi* or with *AkhR* mutation. **Supplementary Video 10**: Ca^2+^ activities in isolated fat body from WT and *Inx2*-*RNAi* adult flies. **Supplementary Video 11**: Ca^2+^ activities in immobilized control, *Inx2-RNAi*, and *AkhR* mutant adult flies.

## Author Contributions

Conceptualization, experiments design by L.H. and N.P.; experiments performance, data collection and analysis by M.A., C.Z. and X.G. All authors have read and agreed to the published version of the manuscript.

## Data Availability Statement

The data presented in this study are available on request from the corresponding author.

## Acknowledgments

We thank Dr. Dalibor Kodríka for sharing the anti-AKH antibody and Dr. Stephanie Mohr for comments on the manuscript.

## Conflicts of Interest

The authors declare no conflict of interest.

This article is subject to HHMI’s Open Access to Publications policy. HHMI lab heads have previously granted a nonexclusive CC BY 4.0 license to the public and a sublicensable license to HHMI in their research articles. Pursuant to those licenses, the author-accepted manuscript of this article can be made freely available under a CC BY 4.0 license immediately upon publication.

---

**Supplementary Table 1:** Fly stocks used in this study.

| Genotypes | Source | Identifier |
| --- | --- | --- |
| *tub-GAL80<sup>TS</sup> ; Lpp-gal4* | Lab Stock |  |
| *UAS-Kir2.1* | Bloomington Stock Center | BDSC_6596 |
| *UAS-Akh-RNAi* | Bloomington Stock Center | BDSC_34960 |
| *UAS-GCaMP5G-T2A-mRuby3* | This study | N/A |
| *Akh-gal4 (on 3rd)* | Bloomington Stock Center | BDSC_25684 |
| *Akh-gal4 (on X)* | This study | N/A |
| *Ilp2-gal4* | Bloomington Stock Center | BDSC_37516 |
| *UAS-Gaq-RNAi* | Bloomington Stock Center | BDSC_33765 |
| *AkhR<sup>null</sup>* | Bloomington Stock Center | BDSC_80937 |
| *UAS-Luc-RNAi* | Bloomington Stock Center | BDSC_35788 |
| *UAS-Gaq* | Bloomington Stock Center | BDSC_30784 |
| *UAS-Serca-RNAi* | Bloomington Stock Center | BDSC_25928 |
| *UAS-AkhR-RNAi* | Bloomington Stock Center | BDSC_29577 |
| *UAS-Ogre-RNAi* | Tsinghua University Fly Center | THU-2608 |
| *UAS-Inx2-RNAi* | Tsinghua University Fly Center | THU-2489 |
| *UAS-Inx3-RNAi* | Tsinghua University Fly Center | THU-0527 |
| *UAS-Zpg-RNAi* | Tsinghua University Fly Center | THU-2759 |
| *UAS-Inx5-RNAi* | Tsinghua University Fly Center | THU-2881 |
| *UAS-Inx6-RNAi* | Tsinghua University Fly Center | THU-2265 |
| *UAS-Inx7-RNAi* | Tsinghua University Fly Center | THU-2180 |
| *UAS-ShakB-RNAi* | Tsinghua University Fly Center | THU-2617 |
| *w<sup>1118</sup>* | Lab Stock |  |
| *Canton S* | Lab Stock |  |

**Supplementary Table 2:**
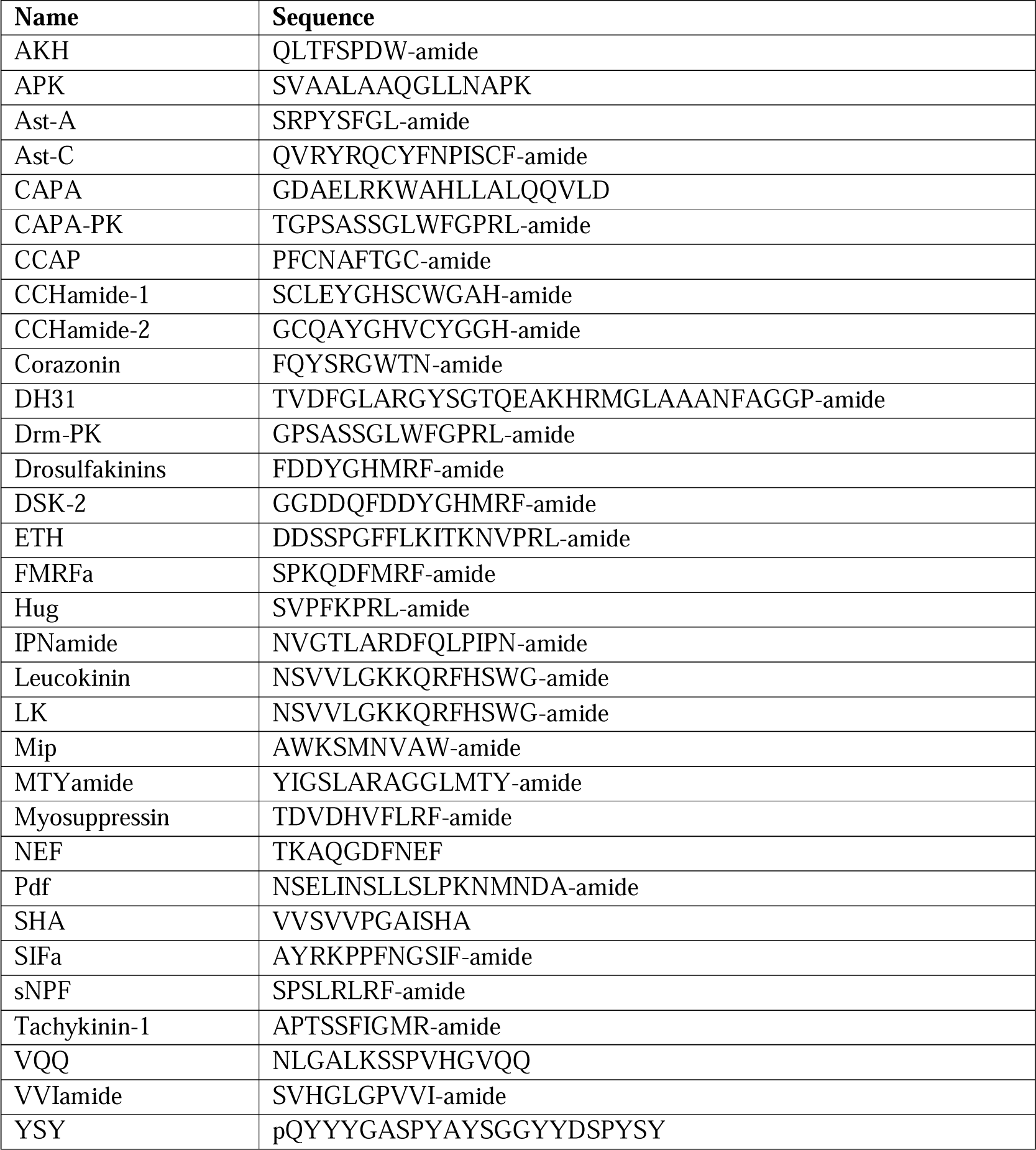
Sequences of neuropeptides used in this study.

**Supplementary Figure 1.**
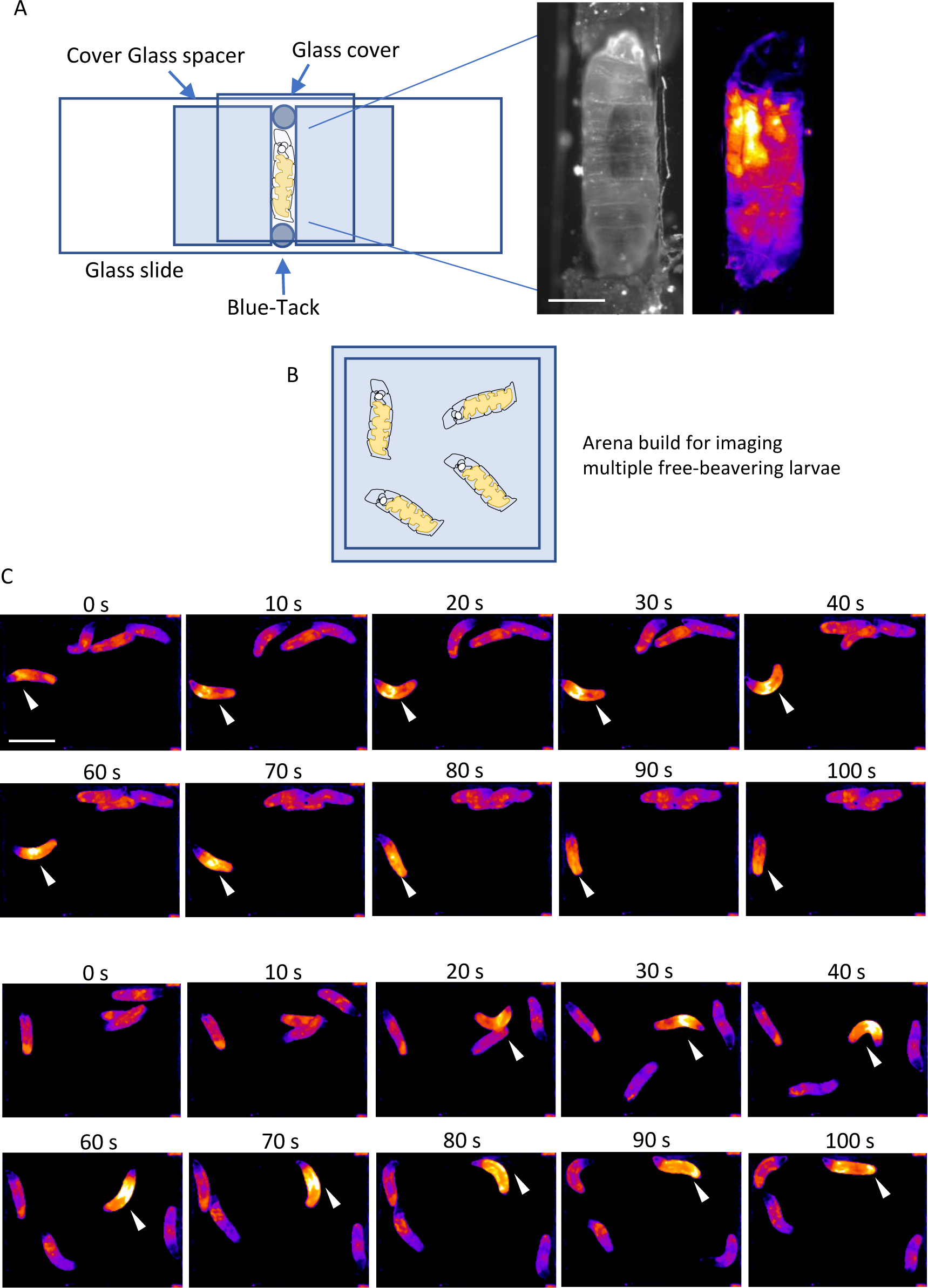
Setups for imaging the *in vivo* Ca^2+^ waves in fly larvae. **(A)** 3^rd^ instar larvae were placed in a narrow glass channel to restrict mobility. Adipose tissue specific Ca^2+^ activity were visualized in *Lpp>UAS-GCaMP5G* animals. **(B)** An arena fitting the imaging field of a fluorescent dissection microscope was generated by 3D-imaging???. A glass was put on the top of the arena to prevent the escape of larvae. **(C)** Ca^2+^ activity in the fat body of free-behaving 3^rd^ instar larvae were revealed in *Lpp-Gal4>GCaMP5G-T2A-mRuby*. Scale bars, 1 mm (A), 5 mm (C).

**Supplementary Figure 2.**
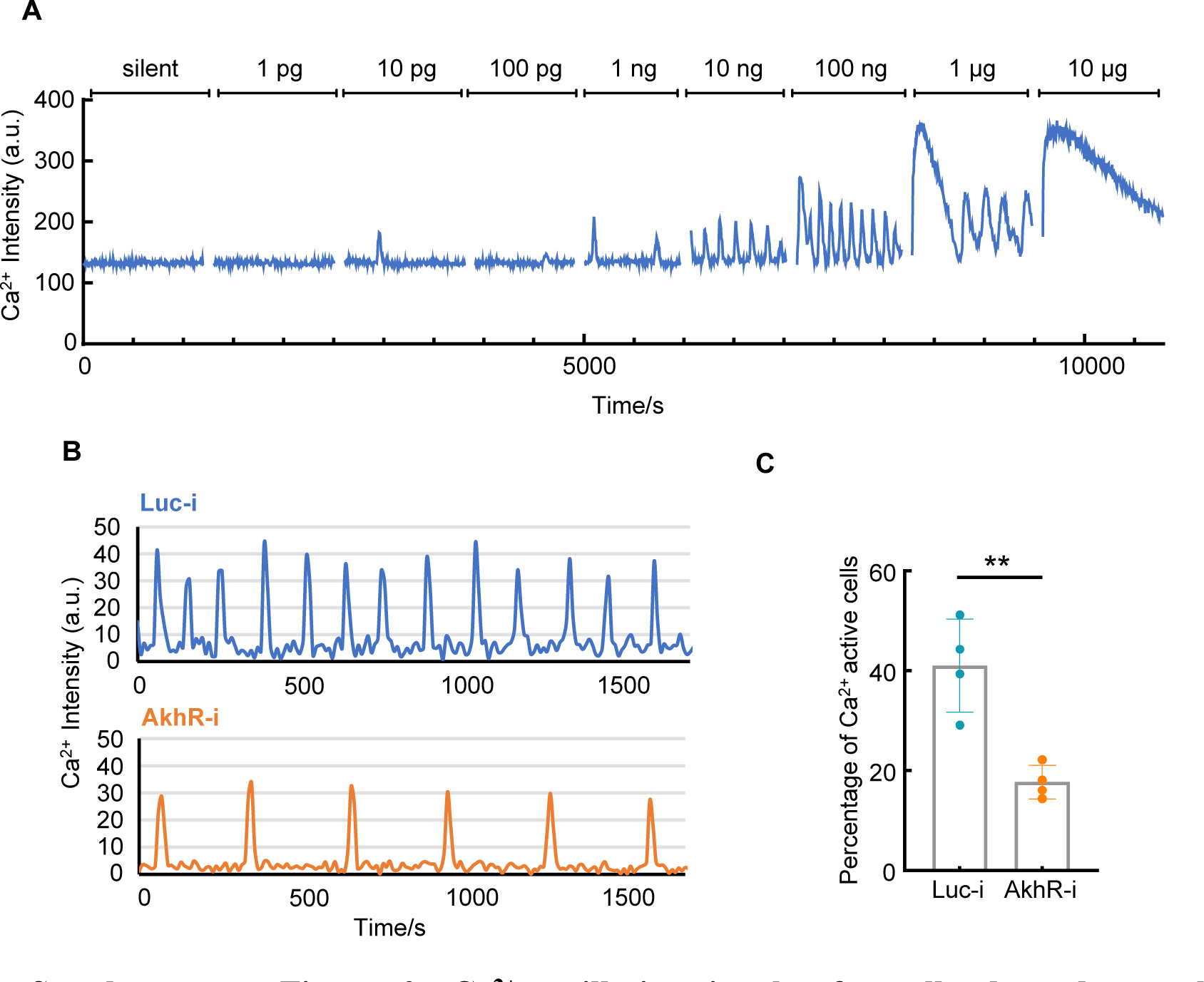
Ca^2+^ oscillation in the fat cells depends on the concentration of applied AKH and the expression level of AkhR. **(A)** 3^rd^ instar larvae expressing *GCaMP5G-T2A-mRuby* were treated with different centration of synthesized AKH ligands. **(B-C)** Fat body expressing *GCaMP5G-T2A-mRuby* together with Ctrl (*luciferase-RNAi*) or *AkhR-RNAi* were treated with 100 ng/mL synthesized AKH peptide. Akh-RNAi significantly reduced the Ca^2+^ oscillation frequency and the percentage of cells with the Ca^2+^ oscillation. Student t-test was used for significant test. Data were plotted as mean ± SD.

**Supplementary Figure 3.**
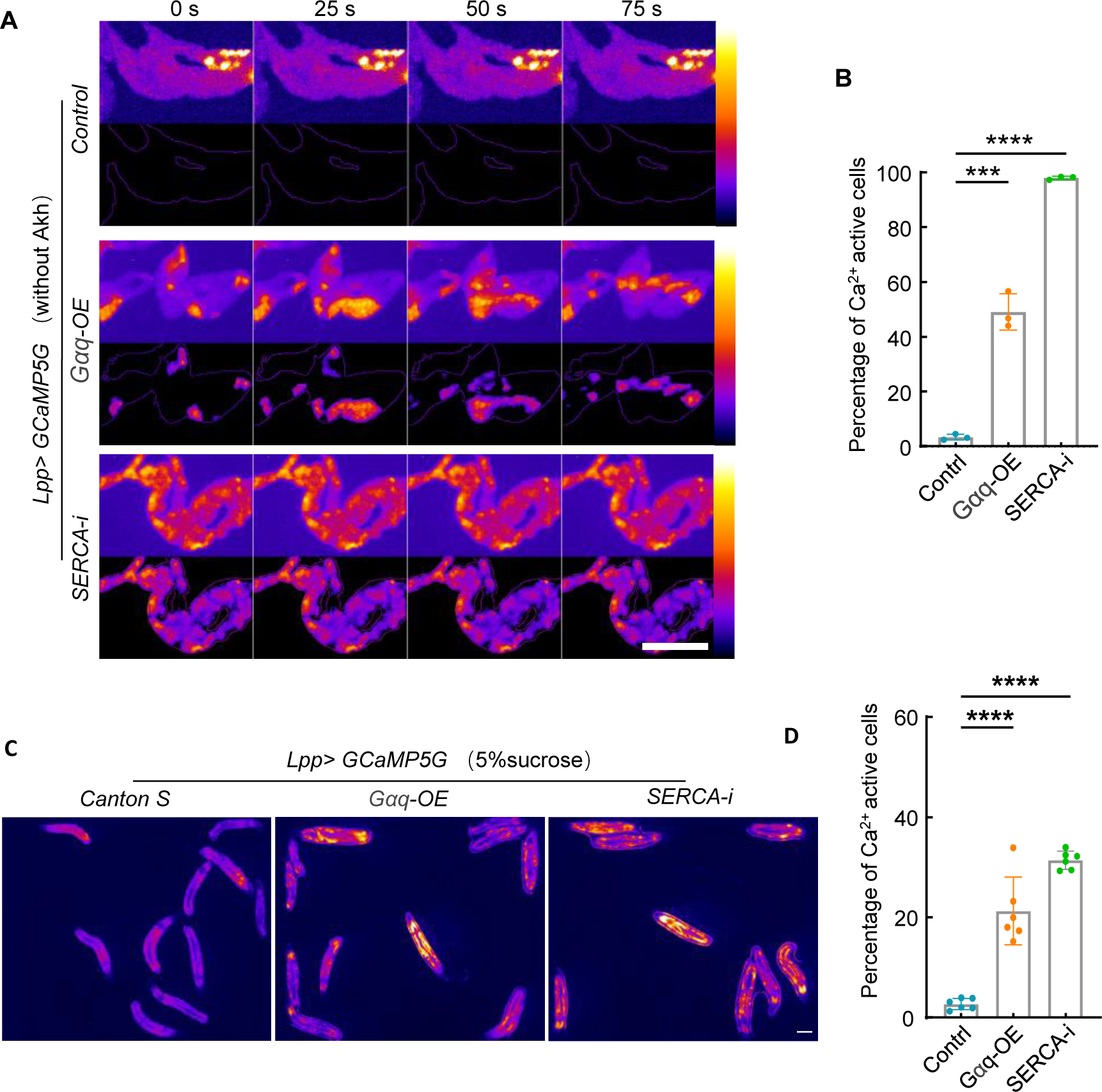
Ca^2+^ activities regulated by Gaq and SERCA in larval fat bodies. **(A-B)** Fat bodies were dissected from 3^rd^ instar larvae expressing GCaMP5G-T2A-mRuby and indicated genes by *Lpp-Gal4*. One-way ANOVA with multiple comparison was used for significant test. **(C-D)** 3^rd^ instar fly larvae expressing *GCaMP5G-T2A-mRuby* and indicated genes by *Lpp-Gal4* were kept on 2% agarose containing 5% sucrose and imaged under free behaving condition. Ca^2+^ activities in the fat bodies were quantified. Data were plotted as mean ± SD. Scale bars, 500 μm (A), 1000 μm (C).

**Supplementary Figure 4.**
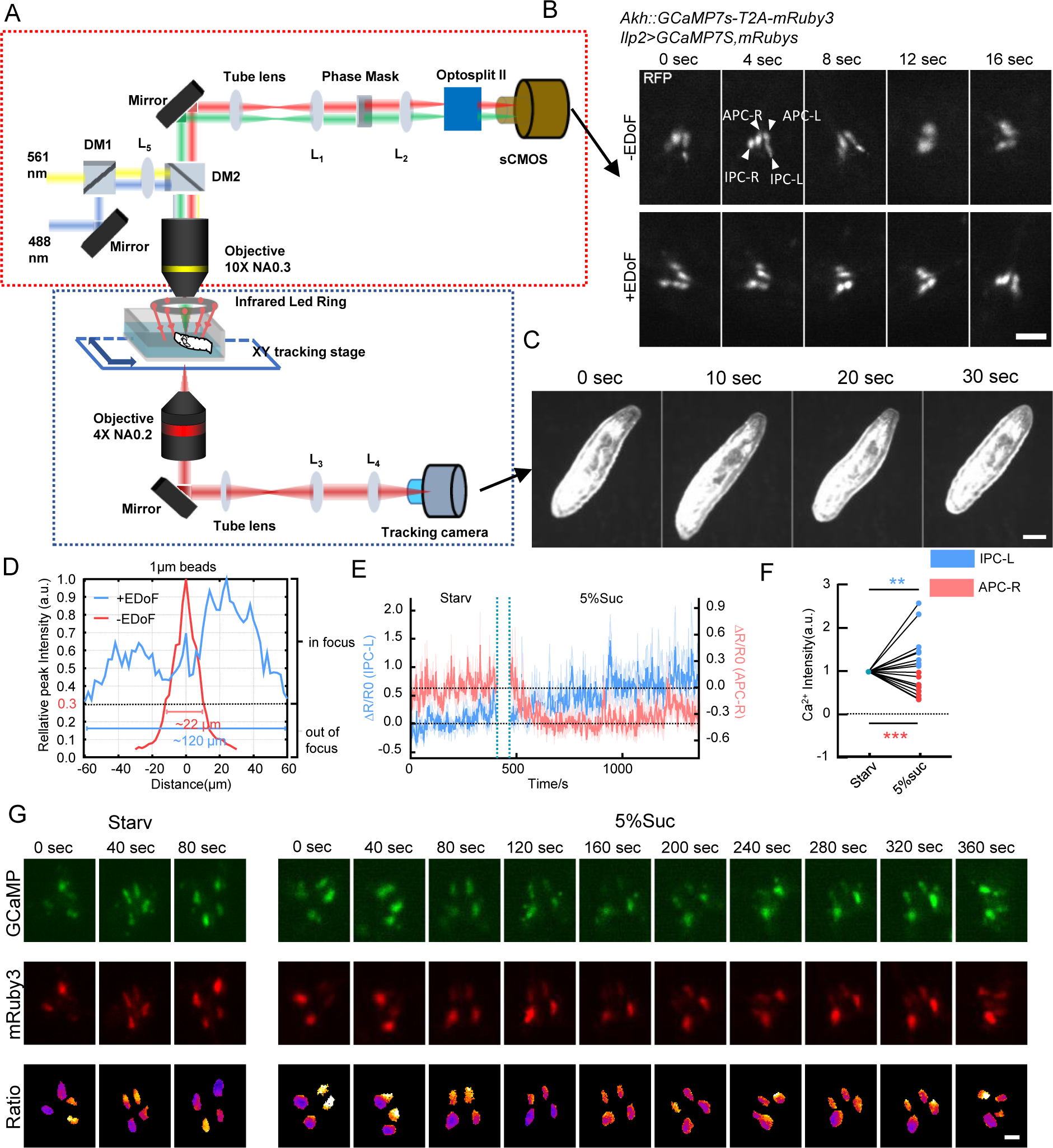
Setups for imaging the Ca^2+^ activities in the APCs of free-behaving fly larvae. **(A)** The Extended-Depth-of-Field (EDoF) microscope has two modules: 1) A darkfield imaging module (blue dotted box) equipped with a 4 x NA 0.2 air objective and a high-speed NIR camera to track and record free-behaving 1st instar larvae **(C)**; and 2) A fluorescence imaging module (red dotted box) equipped with a 10X NA 0.3 air objective and an sCMOS camera, whose imaging surface is bisected by Optosplit II that allows simultaneous recording of two fluorescence signals (calcium-sensitive GCaMP and calcium-insensitive RFP as reference). The microscope’s aspheric phase mask is placed on the conjugate plane of the pupil plane of the objective, i.e., the back focal plane of the achromatic lens. In this way, the point spread function of the system is modified to extend the effective depth of field (we set the out-of-focus standard as a relative peak intensity of 0.3 for 1 μm fluorescent beads) by about 5 times **(D)**. **(B)** Representative RFP images of the APCs and IPCs of free-behaving larvae between –EDoF and +EDoF. After using EDoF technology, out-of-focus situations were reduced. **(E)** Representative Ca^2+^ activities in the APCs (red) and IPCs (blue) of free-behaving larvae transferred between starvation and 5%sucrose. **(F)** Quantification of Ca^2+^ activities in the APCs of free behaving larvae. **(G)** Representative Ca2+ images of the APCs of free-behaving larvae. Scale bars, 40 μm (B), 200 μm (C), 20 μm (G).

**Supplementary Figure 5.**
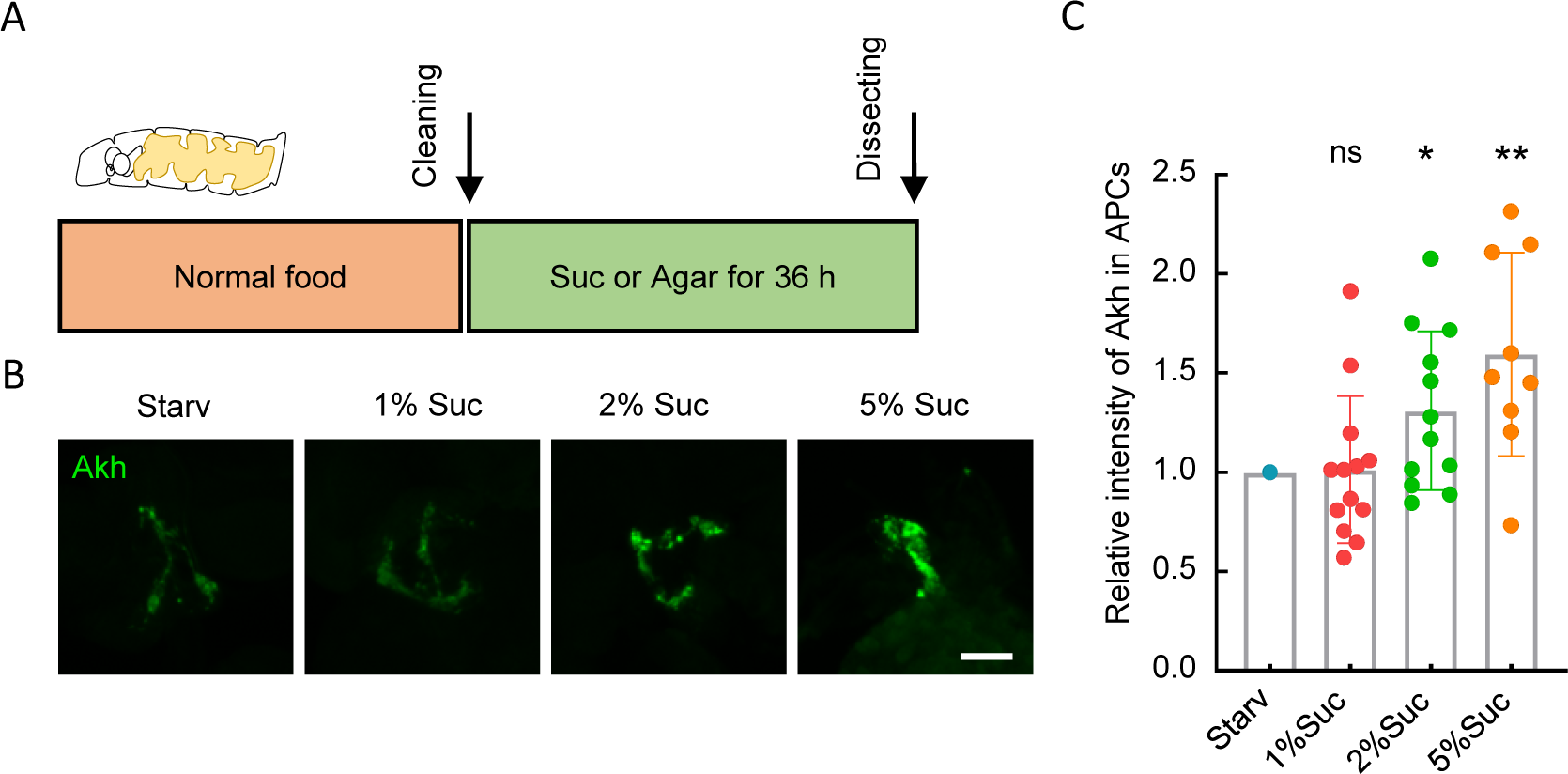
AKH is released after starvation. **(A)** Early 3^rd^ instar larvae were kept on 2% agarose (Starvation) or 2% agarose + different centration of sucrose (Suc) for 36 hours. (B) The remaining AKH in the APCs of the larvae were stained with anti-AKH antibody. **(C)** Quantification of AKH signals in the APCs. Data were plotted as mean ± SD. Scale bars, 25 μm (B).

**Supplementary Figure 6.**
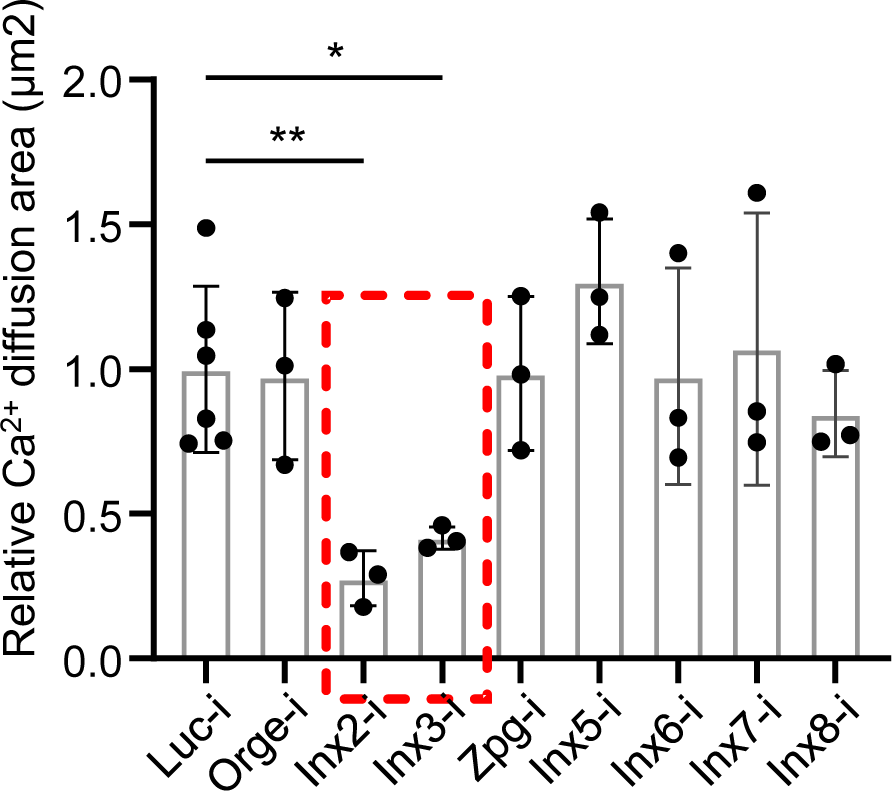
Relative Ca^2+^ diffusion area after knocking down different gap junction proteins in the fly fat body. 3^rd^ instar larvae expressing *GCaMP5G-T2A-mRuby* and indicated RNAi by *Lpp-Gal4* were kept at RT before dissection. Dissected fat bodies were treated with 100 ng/mL synthesized AKH peptide and the Ca^2+^ diffusion areas were quantified as described in the section. Only *Inx2* and *Inx3* knockdown significantly reduced the intercellular Ca^2+^ diffusion. One-way ANOVA with multiple comparison was used for significant test. Data were plotted as mean ± SD.

**Supplementary Figure 7.**
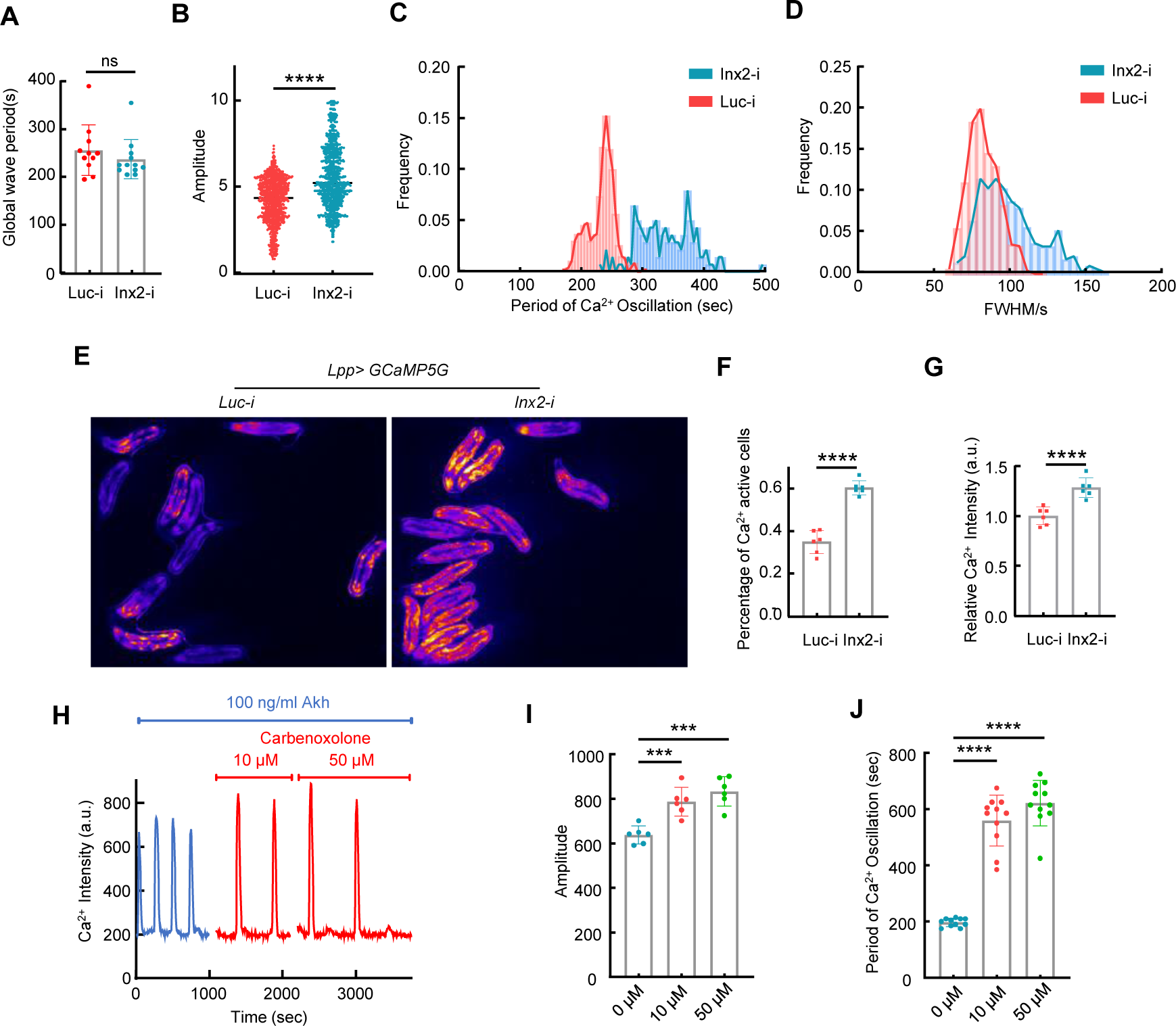
Ca^2+^ activities in the larval fat body is affected by gap junction disruption. **(A)** The period of global Ca^2+^ waves was not affected by *Inx2* RNAi. **(B-D)** Parameters of Ca^2+^ oscillation in *ex vivo* cultured larval fat body activated by 100 ng/mL AKH was quantified. The amplitude, period and peak width (full width at half maxima/FWHM) of the Ca^2+^ signal in the cells increased. **(E-G)** Ca^2+^ activities in the free-behaving WT and *Inx2* RNAi larvae were quantified. Both the proportion of cells with positive Ca^2+^ activities and the average intensity of Ca^2+^ in these cells increased in the larvae with *Inx2* knocked down. **(H-J)** Gap junction inhibitor carbenoxolone triggered similar changes in the amplitude and period of Ca^2+^ waves, resembling those observed upon *Inx2* knockdown. Data were plotted as mean ± SD.

